# Scratching on French PDO cheese surfaces sheds light on an unexplored microbial genomic and metabolic diversity

**DOI:** 10.64898/2025.12.15.694182

**Authors:** Hélène Gardon, Sibylle Tabuteau, Françoise Irlinger, Eric Dugat-Bony, Valérie Barbe, Cécile Callon, Julia Cantuti Gendre, Corinne Cruaud, Céline Delbès, Frédérick Gavory, Valentin Loux, Nacer Mohellibi, Cécile Neuvéglise, Pierre Renault, Olivier Rué, Sébastien Theil, Jean-Marc Aury, Vincent Hervé

## Abstract

Cheeses are fermented dairy products consumed worldwide. Their global diversity results from various local variables, including technological practices, as well as the metabolic activity of diverse microorganisms. In Europe, this typicity is exemplified by Protected Designation of Origin (PDO) cheeses, for which genetic diversity remains largely unexplored. Combining culturomics (*n* = 373 bacterial genomes) and metagenomic (*n* = 146 metagenomes), we performed a national-scale survey of the microbial diversity encompassing 44 French PDO cheeses. Taxonomic (bacteria, fungi and viruses) and functional profiling reveal a high diversity in the cheese rind, mainly driven by the cheese technology. We also reconstructed 1,119 bacterial metagenome-assembled genomes (MAGs) encompassing seven phyla, including *Actinomycetota*, *Bacillota*, *Pseudomonadota* and *Bacteroidota*. Using GTDB as a reference, we identified 221 MAGs encompassing 46 genera, as well as 44 bacterial isolate genomes encompassing eight genera, which represent potentially 81 new species (based on <95% ANI). These species were particularly numerous among the genera *Halomonas*, *Psychrobacter* and *Brachybacterium*. Similar results were observed when compared with the cFMD database. We combined our genomic and metagenomic datasets into a catalog of 26.2 million protein clusters, with 50% of these clusters remaining unassigned to a known function and taxonomy. We illustrated the potential of this resource by searching for methionine gamma-lyase (MGL), an enzyme playing a significant role in cheese flavor. This protein was predominantly found in *Pseudoalteromonas*, a potentially new MGL-producing genus, *Serratia, Pseudomonas, Proteus* and *Hafnia*, and its prevalence varied with cheese technology. Our study provides a substantial genomic resource for food microbiologists and cheesemakers to further explore the biotechnological potential of PDO cheese biodiversity.

## Introduction

Over the last decade, deep metagenomic shotgun sequencing and genome sequencing have allowed the description of most habitats through the creation of microbial gene-^1^ and genome-catalogs ^2–4^. Among these genomes, a large proportion are metagenome-assembled genomes (MAGs) ^5^. These approaches provide the opportunity to explore both the taxonomic diversity and the predicted functions of microorganisms at the genome-level. Besides host-associated and environmental habitats, foods and more particularly fermented foods have been investigated with similar approaches ^6^. A key finding of these global microbial surveys is that many microorganisms remain still undescribed ^6^ and that many genes and functions are rare and habitat-specific ^1^. Therefore, there is a need to explore specific habitats in more detail to reveal their microbial taxonomic novelty and functional potential.

Fermented foods, including cheeses, are consumed worldwide ^7,8^. The broad cheese diversity relies on historical, socio-cultural and technological practices as well as on a wide microbial diversity ^9–12^. Indeed, hundreds of cheese varieties have been described ^13^, resulting from various technologies used in cheese production (e.g. hard, semihard, soft, and fresh cheese) ^14^. In Europe, some of this diversity can be found in cheeses with a Protected Designation of Origin (PDO) label. These cheeses are produced, processed and made in a defined geographical area according to strict specifications and traditional know-how. They are widely acknowledged for their high quality and non-standardised nature.

Cheese microbial diversity encompasses both microbial starters added by the cheesemaker and endogenous microbes naturally present in the milk and the surrounding environment. Taxonomically, they include bacteria, fungi and viruses ^12,15,16^. Over the years, the taxonomic diversity of these cheese microbes has been largely investigated using both culture-dependent ^17,18^ and culture-independent methods such as metabarcoding ^19–22^. These numerous metabarcoding studies have improved our understanding of the assembly rules of these microbial communities. These communities result from a combination of processes ^23^ with both biotic and abiotic factors being essential drivers of the community assembly ^21,22,24–26^. Indeed, salinity, pH, humidity and temperature can significantly shape the cheese microbiota and these variables are also key features of certain cheese families, as well as technological practices imposed by PDO cheese specifications. However, metabarcoding approaches show limited resolution; hence, there is a need for genome-scale resolution to explore hidden microbial diversity and predict functions ^27,28^. To date, a few PDO cheeses have been investigated with shotgun metagenomics ^29–31^ to assess cheese quality and safety. Additionally, comparisons of bacterial genomes from PDO cheeses revealed genetic diversity and specificity, but such studies have so far focused on lactic acid bacteria ^29,32,33^ while the phylogenetically diverse ripening cheese bacteria have been overlooked.

Recently, Irlinger et al ^22^ highlighted the importance of both geographical and human factors in the assembly of microbial communities of French PDO cheeses, using a massive metabarcoding dataset. The present study, built on a subset of the same samples, follows-up on Irlinger study by combining shotgun metagenomics and bacterial culturomics to study samples from 44 PDO cheeses present across France and belonging to 7 different cheese-making technologies. In the present study, we aimed to *i*) explore at the national-scale the taxonomic and functional diversity of microorganisms (fungi, bacteria and viruses) associated with French PDO cheeses; *ii*) investigate the impact of cheese technology on the cheese microbiome diversity; *iii*) establish a large genomics and metagenomics resource to answer both fundamental and applied research questions.

## Materials and Methods

### Cheese sampling and bacterial isolation

Cheese samples were collected for metagenomic analysis from both the rind and core cheese, as well as from milk, following the procedure described in ^22^. In parallel, we isolated 373 bacterial strains from 117 French PDO cheeses produced in various regions (Irlinger et al., 2024). One g from each sample was taken and homogenized in 10 mL sterile saline solution (9 g/l NaCl) with an Ultra Turrax (VWR International, Fontenay-sous-Bois, France) at 8,000 rpm for 1 min). The mixture was then serially diluted in 10-fold steps in sterile saline solution (9 g/l NaCl) and plated on i) Brain Heart Infusion (BHI) Agar (Difco, Detroit, MI, USA) supplemented with 44 mg/l amphotericin B for the isolation of aerobic bacteria, incubated at 28°C; ii) Marine Agar in which we added from 0 to 4% salt (w/v) for the isolation of halotolerant and halophilic bacteria, incubated at 15°C and 28°C for 3 to 5 days. After the incubation period, twenty dominant bacterial colonies were randomly selected from countable plates from each cheese sample and were purified by streaking twice on the appropriate media. The purified strains were cultured in 10 ml appropriate medium for 24 hours with continuous agitation. Cell cultures were either harvested by centrifugation and pellets frozen for later extraction of genomic DNA or directly stored at −80°C in a 1:1 mixture of glycerol and appropriate medium.

### DNA extraction and sequencing

#### Metagenomic DNA extraction

A 250-mg aliquot of cheese samples was added to a 2-ml tube containing 350 mg of zirconium beads (diameter, 0.1 mm; Sigma, St-Quentin-Fallavier, France), 250 µl of guanidine thiocyanate (4 M) in Tris-HCl (0.1 M, pH 7.8) and 40 μl N-lauryl sarcosine (10%) and shaken in a Precellys Evolution bead beater (Bertin, Montigny-le-Bretonneux, France) for 20 sec at a speed of 6 500 m/s. 75 μl of lysozyme (3 mg) and lyticase (100 unity) were added, and the tube was incubated for 30 min in a water bath at 37°C. 40 μl of proteinase K (15 mg/ml) and 100 μl of sodium dodecyl sulfate (20%) were then added. The tube was incubated for 30 min in a water bath at 55°C. 200 μl of sodium phosphate buffer (0.1 M, pH 8), 200 μl of 50 mM acetate-10 mM EDTA buffer (pH 5), and 500 μl of phenol-chloroform (25:24:1, pH 8) were then added. The tube was vigorously shaken in a Precellys Evolution bead beater (Bertin, Montigny-le-Bretonneux, France) for 45 sec at 10000 m/s. The tube was incubated for 2 min at 55°C and cooled on ice for 5 min. After another 45 seconds of mixing 10000 m/s), the tube was incubated for 2 min at 70°C, cooled on ice for 5 min, and centrifuged for 20 min at 14,000 rpm and room temperature, separating two phases. The upper aqueous phase was recovered in a 2-ml tube (Phase Lock Gel Heavy; Eppendorf, Hamburg, Germany) and mixed with 500 µl of phenol-chloroform (25:24:1, pH 8). The tube was centrifuged for 5 min at 14,000× g and 20°C. Chloroform (500 µl) was then added to the tube, which was mixed gently. After a third centrifugation, the aqueous phase was recovered, mixed with two μl of RNase A (20 mg/ml; SERVA Electrophoresis GmbH, Heidelberg, Germany), and incubated for 30 min at 37°C. The extracted DNA was then purified and concentrated with the Genomic DNA Clean & Concentrator-10 kit (Zymo Research). A blank extraction (adding nothing to the bead tube) was performed alongside sample extractions for the negative extraction controls. DNA extracts were stored at −20 °C until further analysis.

To extract DNA from milk, we thawed 180 mL samples in a water bath at 25°C, then 12 mL of SDS (sodium dodecyl sulfate 20% solution) were added to 120 mL of milk, which were then incubated for 30 min at 30°C. After centrifugation (5300× g, 30 min, 4 ◦C), the fat layer and the supernatant were removed. The pellets obtained were mixed with 1 mL sterile PBS (Phosphate-buffered saline), incubated for 10 min at 30°C, centrifuged (13,000× g, 5 min, 4°C) and stored at −20°C. Total DNA extraction was then performed as described for cheese samples.

#### Genomic DNA extraction for short-read sequencing

The total genomic DNA of 373 strains was extracted using the Nucleospin Tissue kit (Macherey-Nagel, Dueren, Germany) according to the manufacturer’s instructions with the additional step of incubating the previously frozen bacterial pellets in 180 μl lysis buffer [2 mM EDTA, 20 mM Tris (pH = 8.0), lysozyme (20 mg/ml), mutanolysin (10 units) and 1.2% Triton® X-100] for 30 minutes at 37°C, prior to the enzymatic lysis step (200 µl of 20 mg/ml proteinase K and B3 lysis buffer at 56°C for 30 min and RNase treatment (10 mg/ml). The other steps were then performed according to the Nucleospin Tissue Kit manual (Macherey-Nagel, Dueren, Germany).

#### Genomic DNA extraction for long-read sequencing

The total genomic DNA of 36 bacterial strains was extracted using the NucleoBond HMW DNA kit (Macherey-Nagel, Düren, Germany) according to manufacturer’s protocol with the additional step of incubating for 30 minutes at 37°C, the previously frozen bacterial suspension (100 mg in 500 µl of water) with 500 µl lysis buffer [2 mM EDTA, 20 mM Tris (pH = 8.0), lysozyme (20 mg/ml), mutanolysin (10 units) and 1.2% Triton® X-100] prior to the enzymatic lysis step (200 µl of 20 mg/ml proteinase K and 200µl of H1 lysis buffer at 50°C for 30 min and RNase treatment (10 mg/ml). The other steps were then performed according to the NucleoBond HMW DNA Kit manual (Macherey-Nagel, Dueren, Germany).

#### Sequencing

Both metagenomes and bacterial genomes were sequenced on Illumina NovaSeq 6000 (2×150 bp). Additionally, 36 bacterial strains were also sequenced with Oxford Nanopore Technologies (PromethION and GridION Mk1) to perform hybrid assembly.

### Sequence processing

For both genomes and metagenomes, short Illumina reads were bioinformatically post-processed *sensu* Alberti et al ^34^ to filter out low quality data. First, low-quality nucleotides (Q < 20) were discarded from both read ends. Then, the remaining Illumina sequencing adapters and primer sequences were removed and only reads ≥30 nucleotides were retained. These filtering steps were done using in-house-designed software based on the FastX package (https://www.genoscope.cns.fr/fastxtend/). Finally, read pairs mapping to the phage phiX genome were identified and discarded using SOAP aligner^35^ (default parameters) and the *Enterobacteria* phage PhiX174 reference sequence (GenBank: NC_001422.1).

For bacterial genomes, Illumina reads were processed with fastp v0.23.1 ^36^ and *de novo* assembly was performed with Unicycler v0.4.8 ^37^ (mode normal). For bacterial genomes sequenced with both technologies, a hybrid assembly method was applied using short– and long-reads with Unicycler v0.4.8. Quality assessment was done with Quast v5.0.2^38^.

To eliminate potential host contamination, metagenomic reads were aligned against the genome of the animal from which the cheese milk is derived [*Bos taurus (*AC_000158.1), *Capris hircus* (NC_030808.1) and *Ovis aries* (NC_019458.2)] using bwa v0.7.17 ^39^. The quality controlled metagenomic reads were then assembled into scaffolds using SPAdes v3.14.0 ^40^ (*meta* option for metagenomic datasets). Quality metrics and statistics of each metagenome assembly were evaluated with metaQUAST v5.0.2 ^41^. Bacterial metagenomes-assembled genomes (MAGs) were reconstructed within each metagenome with *SnakeMAGs* workflow v1.0.0 ^42^, from the binning of the scaffolds with MetaBAT2 ^43^ to the estimation of the relative abundance of MAGs.

MAGs and genomes were quality controlled with CheckM v1.1.3 and GUNC v1.0.5 ^44^. Only genomic sequences with completeness ≥ 50%, contamination <10% and treated as non-chimeric were kept. The taxonomic assignment was done using GTDB-Tk v2.1.0^45^ using the Genome Taxonomy Database (GTDB) r207 ^46^. We also used dRep to cluster all the MAGs and genomes not assigned to the species level (<95% ANI) and thus quantify the number of potentially new species clusters ^47^.

For metagenomic scaffolds, genomes and MAGs, protein-coding sequences (CDSs) were predicted with Prokka v1.14.6 ^48^ (with the parameter ‘––metagenome’ for scaffolds to improve gene predictions). CDS were further annotated with eggNOG-mapper v2.1.8 ^49^ and the EggNOG database 5.0.2 ^50^.

### Taxonomic profiling and phylogenomics

We performed a metagenomic taxonomic profiling of the 146 metagenomes. Prokaryotic and eukaryotic compositions and abundances were obtained using mOTUs v3.0.3 ^51^ and EukDetect v1.3 ^52^ respectively. For the viral composition, we first used VIBRANT v1.2.1 ^53^ on scaffolds ≥2kb to identify viral metagenomic scaffolds, and subsequent low quality drafts were removed. CheckV v0.8.1 ^54^ was used on viral sequences to assess their completeness and remove flanking host regions on assembled proviruses. Only genomes with >50% estimated completeness were selected and then clustered into viral operational taxonomic units (vOTUs) using 95% average nucleotide identity (ANI) over 85% of the length of the shorter sequence ^55^ with MMseqs2 v14.7e284 ^56^ and the following parameters: ––cov-mode 1, ––cluster-mode 2, ––min-seq-id 0.95, –c 0.85. vOTUs abundance was estimated by mapping metagenomic reads on the vOTU sequences with bwa v0.7.17 ^39^. Taxonomic assignment of the vOTUs was performed with geNomad v1.5.0 and the associated database v1.2 ^57^. Finally, vOTUs sequences were compared by Blastn v2.16.0 ^58^ to an in-house database consisting of genome sequences from 32 common dairy phages (available here: https://forge.inrae.fr/eric.dugat-bony/cheese_virome/-/tree/91bc85bcb7215b79fc4a583178ec 1d5eb1b76e3d/Dairy_phages_database) to identify potentially related phages with known taxonomy and verified host. Based on these analyses, we considered the results of mOTUs, EukDetect and vOTUs as proxy for bacterial, eukaryotic and viral species. The hierarchical taxonomy was visualized with Graphlan v1.1.3 ^59^.

Bacterial genomes and MAGs from this study were used to reconstruct the phylogenomic tree based on 120 aligned and concatenated genes extracted with GTDB-tk v2.4.0. A maximum likelihood tree was inferred with IQ-TREE v2.1.4 ^60^ using the LG+F+I+G4 model of amino-acid evolution (based on ModelFinder selection). The resulting tree was visualized with R package *ggtree* ^61^. To compare the degree of novelty among the bacterial genomes (MAGs and genomes from strains) generated in the present study, we computed with fastANI v1.34 ^62^ the average nucleotide identity (ANI) between our genomes and the cFMD database v1.1.0 ^6^. To date, this database represents the largest genomic resource for food, with 10,112 prokaryotic MAGs reconstructed from 2,533 food-associated metagenomes. We used the standard ANI threshold of 95% ANI cutoff for species demarcation ^62^.

### Protein catalog

We used the protein-level assembly software PLASS v4.687d7 ^63^ to build a protein catalog from the QC metagenomic reads. Then we also collected all the protein sequences from the predicted from the vOTUs, the bacterial MAGs and genomes of bacterial strains and clustered them with the PLASS catalog using the easy-linclust algorithm of MMseqs2 v14.7e284 ^64^ with the following parameters: ––cov-mode 1 –c 0.8 ––cluster-mode 2 ––min-seq-id 0.90. We estimated the number of complete proteins based on the presence of both start and stop codon in a sequence. We used the *rarefaction.single* function from *mothur* v1.48.0^65^ to generate rarefaction curves. The whole catalog was annotated taxonomically using MMseqs2 with UniRef100 database (release-2023_03) ^66^, and functionally with eggNOG-mapper v2.1.8 ^49^ and the EggNOG database 5.0.2 ^50^.

### Methionine gamma–lyase detection and classification

To search for the presence of methionine gamma-lyase in our catalog, we use an HMM profile with HMMER v3.3.2 ^67^. We built a custom profile using a set of manually curated sequences extracted from InterPro on May 14^th^, 2024 ^68^. First, 2,502 sequences were clustered at 95% identity and coverage with MMseqs2. Then, the resulting 1,251 sequence clusters were aligned with MAFFT v7.487 ^69^ and used to generate the profile with *hmmbuild.* The catalog was searched with *hmmscan* (e-value 0.00001) and only sequences longer than 349 AA, shorter than 751 and with a bit-score of minimum 550 were retained. Each sequence was taxonomically assigned using genome/MAG taxonomy. Furthermore, this classification was validated by a phylogenetic approach. Both curated reference sequences and sequences from our catalog were aligned with MAFFT v7.487 ^69^ and used to infer a maximum likelihood tree with IQ-TREE v2.1.4 ^60^.

### Diversity and statistical analyses

To estimate the coverage and sequence diversity of each metagenomic sample, we applied Nonpareil v3.4.1 with the k-mer algorithm (*k* = 32) on the forward reads ^70^. All statistical analyses were performed with R version 4.2. Data manipulation and visualization was done with *tidyverse* collection of R packages ^71^. Multiple comparisons between environments or technologies were performed with Kruskal-Wallis tests followed by Dunn tests with Bonferroni correction. Ordination (NMDS) and PERMANOVA (adonis2) were performed with the *vegan* package. To identify COGs with differential abundance between technologies, we used the ALDEx2 algorithm ^72^. In this analysis, statistical significance was assessed based on the Kruskal-Wallis test with Benjamini and Hochberg’s correction to maintain a 5% false discovery rate. The abundance of these COGs was visualized with the *ComplexHeatmap* package ^73^.

## Results

### Overview of the metagenomic dataset

Our national-scale metagenomic survey comprised 146 samples (*n* = 115 cheese rind, *n* = 22 cheese core and *n* = 9 milk samples) encompassing 44 PDO cheeses present across France and belonging to 7 different cheese-making technologies. After quality filtering, a total of 2.26 billion reads (ca. 341 Gbp) were generated with a median value of 15.7 million reads (ca. 2.4 Gbp) per sample (Table S1). Coverage estimation was high for all samples (min = 90.4%; max = 99.8%; median = 98.7%), indicating that the sequencing effort was appropriate (Figure S1A). We found a significantly higher sequence diversity in rind cheese than in milk and core cheese metagenomes (Figure S1B). Interestingly, we also observed a positive and significant relationship between sequence diversity and pH (r = 0.67, *P* < 0.001) and this relationship remains true when considering core and rind samples separately (Figure S2).

Taxonomic assignment of the unassembled reads revealed that both milk and core samples were dominated by bacterial reads (median 96.3% and 87.7% for milk and core metagenomes, respectively) (Figure 1A). The composition was more variable for the rind samples, with a compositional balance and linear relationship (r = –0.97, *P* < 0.001) between eukaryotic and bacterial reads that reflected the cheese-making technology (Figure 1B and C). Indeed, rind samples from soft bloomy rind, internal blue mold and lactic bloomy rind tended to have significantly higher eukaryotic / bacterial read ratio than rind samples from uncooked pressed cheese, semihard cheese, soft washed rind and hard cooked cheese (*P* < 0.05). Viral sequences were detected in all samples but appear to be more abundant in cheese than in milk. We found no correlation between viral read abundance and bacterial (r = –0.13, *P* = 0.113) or eukaryotic (r = 0.03, *P* = 0.691) read abundance. After manual examination, no archaeal reads nor contigs were identified. Overall, similar taxonomic patterns and conclusions were observed using assembled contigs (Figure S3).

**Figure 1.**
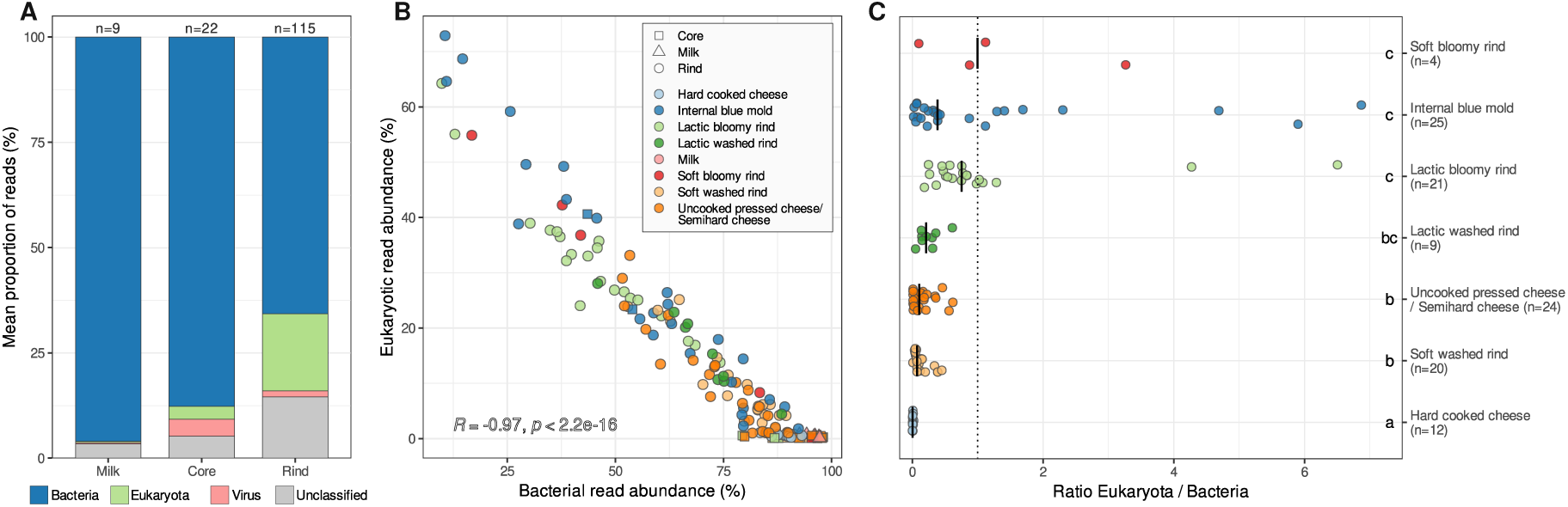
Taxonomic composition of the metagenomic samples based on read classification with Kaiju. **A.** Average taxonomic distribution of the main clades in milk, core and rind cheese samples. **B.** Relationship between eukaryotic and bacterial reads. Pearson correlation coefficients (R) is presented. **C.** Ratio of eukaryotic and bacterial reads in the rind cheese samples.

### Diversity pattern and taxonomic distribution

For each of the main clades (bacteria, eukaryotes, viruses), we used dedicated bioinformatic tools to identify metagenome-based species (see Method section) and investigate taxonomic and diversity patterns. We found that species richness was significantly higher in the rind than in the milk for bacteria, eukaryotes and viruses (*P* < 0.05; Figure 2). When considering only the rind samples, a significant effect of the technology was also observed for the species richness of these three main taxonomic groups (Figure 2A, D, G). We found no correlation between eukaryotic, bacterial or viral richness (*P* > 0.05).

**Figure 2.**
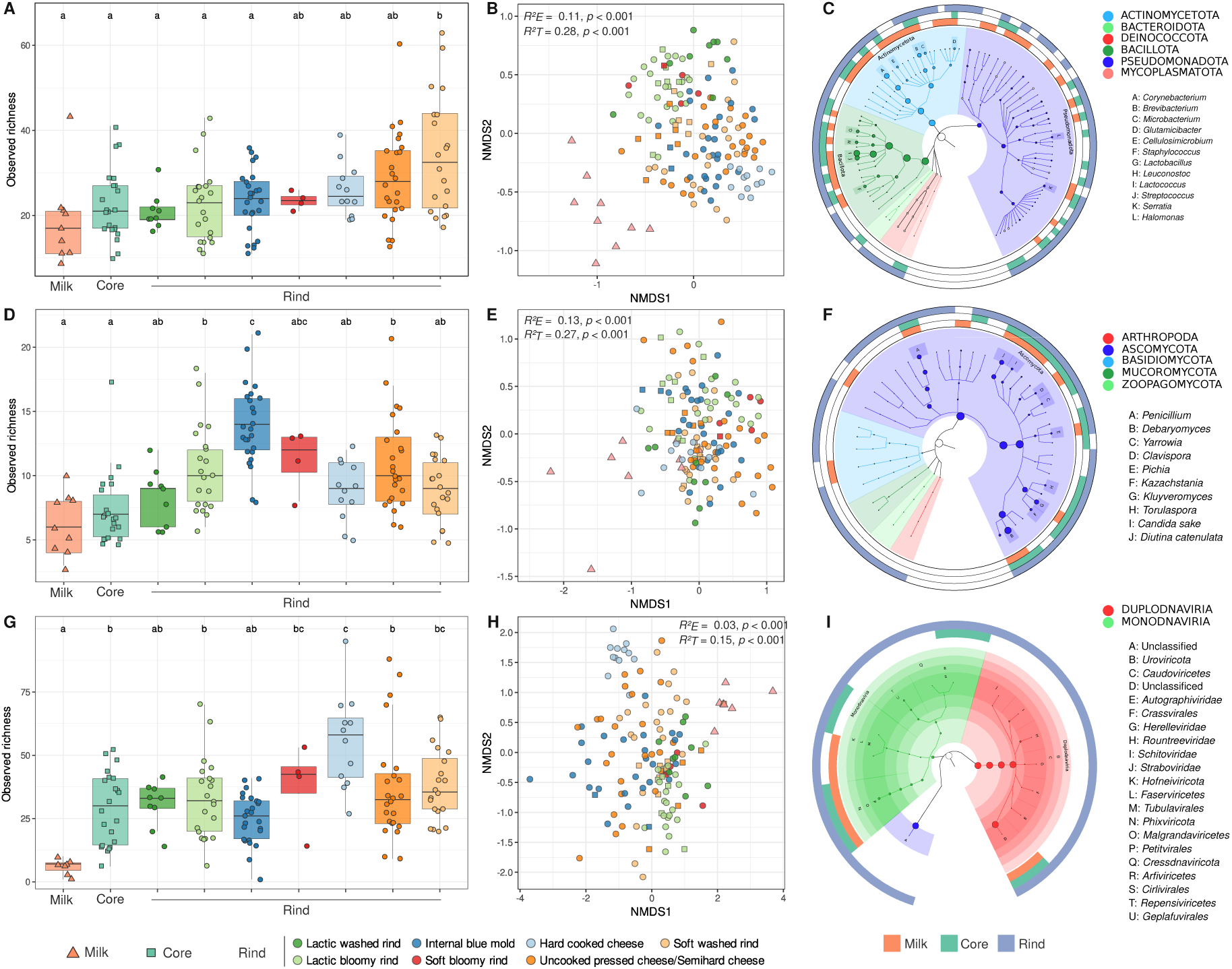
Diversity pattern and taxonomic distribution based on taxonomic profiling of the metagenomic samples. **A.** Bacterial richness. **B.** NMDS ordination of the bacterial community composition based on Bray-Curtis distances. **C.** Taxonomic distribution of the bacteria. **D.** Eukaryotic richness. **E.** NMDS ordination of the eukaryotic community composition based on Bray-Curtis distances. **F.** Taxonomic distribution of the eukaryotes. **G.** Viral richness **H.** NMDS ordination of the viral community composition based on Bray-Curtis distances. **I.** Taxonomic distribution of the viruses. Differences in richness between sample categories were evaluated by a Kruskal-Wallis test followed by post-hoc tests with Bonferroni correction (*P* < 0.05). Effects of the environment (E, milk/core/rind) and cheese-making technology (T, the seven cheese technologies) on the community compositions were evaluated by PERMANOVA.

Regarding the *beta* diversity, community composition significantly differs between milk, core and rind cheese samples for bacteria, eukaryotes and viruses (Figure 2B, E and H; PERMANOVA, *P* < 0.001). Similarly, significant differences were observed when testing the effect of the technological family on the rind samples (PERMANOVA, *P* < 0.001). Rind samples from certain technologies showed particularly high similarity; for instance, hard cooked cheese samples showed clear and distinct clusters especially for bacterial (Figure 2B) and viral (Figure 2H) *beta* diversity. On the contrary, internal blue mold cheese exhibited more heterogeneity with larger and more dissimilar clusters (Figure 2).

These community composition differences were also visible at the taxonomic level (Figure 2C, F and I). Among bacteria, we identified a total of 264 species (mOTUs). As expected, lactic acid bacteria including species of *Lactobacillus, Lactococcus, Streptococcus, Bifidobacterium* and *Leuconostoc* were found in both cheese and milk samples (Figure 2C). On cheese rind, we found 214 bacterial species encompassing 72 genera including well known members of the cheese ripening bacteria, namely *Corynebacterium*, *Brevibacterium*, *Halomonas, Vibrio*, *Microbacterium, Glutamicib*acter, *Pseudoalteromonas*, *Hafnia*, *Propionibacterium*, *Psychrobacter* (Table S2). Among Eukaryota, we identified 77 fungal species encompassing genera traditionally detected in cheese, including *Penicillium, Pichia, Trichosporon, Mucor, Candida, Debaryomyces, Kazachstania, Kluyveromyces* (Figure 2F). Noteworthy, we also recovered sequences belonging to a cheese mite (Arthropoda), namely *Tyrophagus putrescentiae*, in a rind sample of blue cheese (Table S2). Regarding viruses, we identified 976 vOTUs belonging to two main realms, namely *Duplodnaviria* and *Monodnaviria,* as well as 48 unclassified vOTUs (Figure 2I). Using an in-house curated dairy phage database (see Method section), we identified 169 (17%) vOTUs sharing sequence homology with known phages previously isolated from dairy products, including both LAB phages and recently discovered virulent phages infecting ripening bacteria. The most abundant and prevalent vOTUs present in the dataset correspond to *Lactococcus* phages belonging to the well-known 936, P335, C2, and 949 groups.

### Functional diversity

Metagenomic data were also used to predict functions. We found a total of 549,954 clusters of orthologous functions (COGs), with a median of 4,214 COGs per metagenome. Significant differences in COG richness were detected between the different environments and technologies (Figure 3A). In particular, we observed significantly more COGs in rind than in milk and core samples (*P* < 0.05). Among rind samples, hard-cooked cheeses exhibited significantly less COGs than in other technologies (*P* < 0.05).

**Figure 3.**
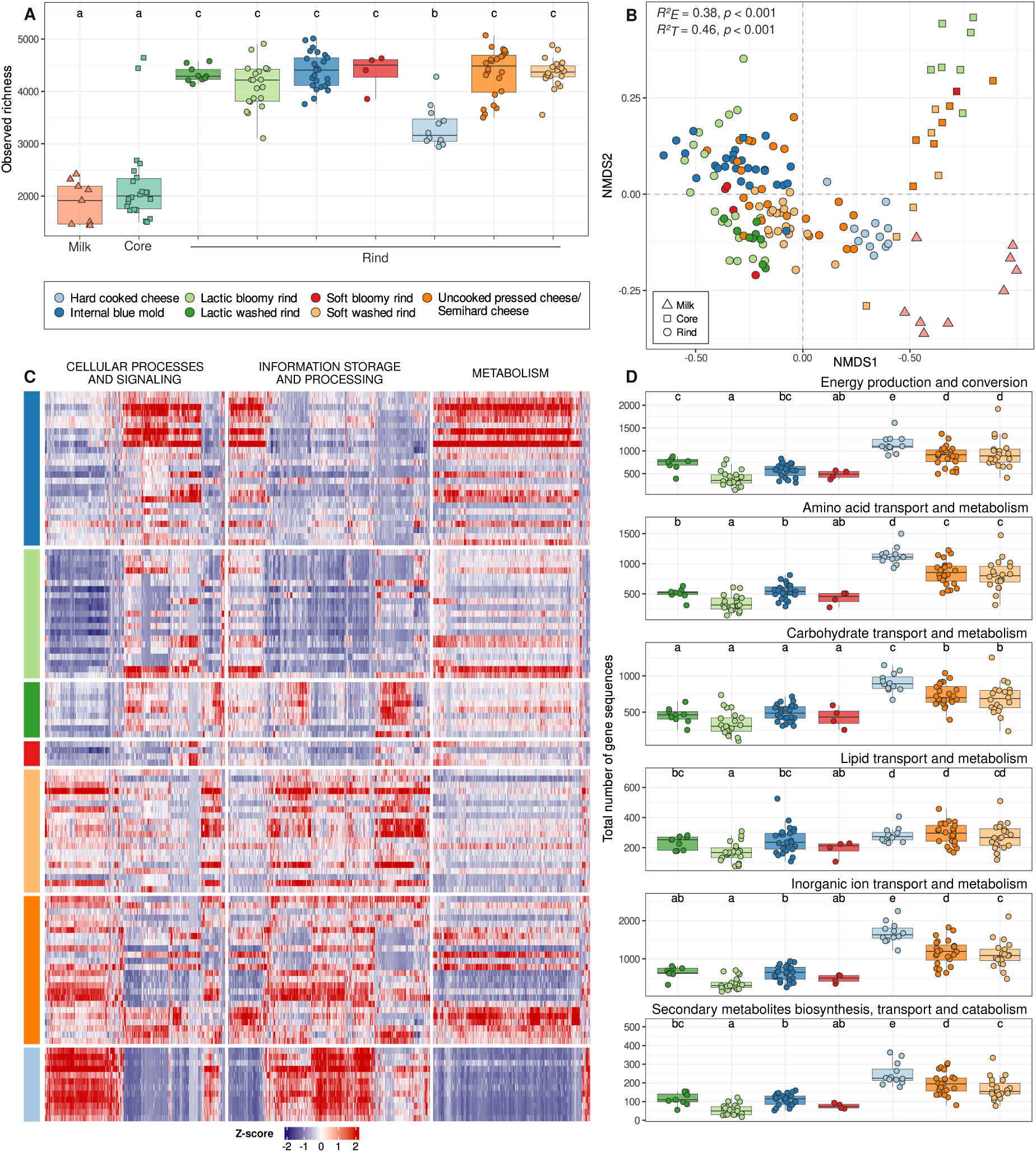
Diversity pattern of the predicted functions from the metagenomes. **A.** Richness of COGs in the different metagenomic samples. **B.** NMDS ordination of the COG composition based on Bray-Curtis distances. **C.** Heatmap showing the differentially abundant rind COGs between cheese technologies (Aldex2 results). Data was scaled with z-score and sorted by cheese technology (rows) and COG categories (columns). **D.** Boxplots showing significant differences in number of gene sequences in rind samples for different COG metabolism categories. Statistical comparisons were computed with Kruskal-Wallis test followed by Dunn’s test with Bonferroni adjustment.

Analysis of functional *beta* diversity revealed significant differences between milk, core and rind, as well as, between the rind samples from different technologies (Figure 3B, PERMANOVA, *P* < 0.001). Interestingly, for both the sample origin and cheese technology factors, more variance was explained by functional than taxonomic *beta* diversity. Again, hard-cooked cheeses formed a coherent and distinct cluster compared to other technologies, such as lactic bloomy rind cheeses and uncooked pressed cheeses showing more dissimilarity (Figure 3B).

Examining the differentially abundant COGs in the rind samples from different cheese technologies, we identified 4,337 significant functions. A heatmap of the COG abundances revealed distinct patterns between the cheese technologies among the main COG categories, namely cellular processes and signalling, information storage and processing, and metabolism (Figure 3C). An in-depth analysis of the latest category revealed that hard-cooked cheese rinds, semihard cheese rinds and soft-washed rinds possessed more gene sequences for energy production and conversion, as well as for the transport and metabolism of amino acids, carbohydrates, inorganic ions and secondary metabolites, compared to other technologies (Figure 3D).

### Genomic diversity

Using the 146 metagenomes, we reconstructed 1,119 bacterial MAGs (>50% completeness, <10% contamination) encompassing seven phyla according to GTDB, namely *Actinomycetota* (*n* = 443), *Bacillota* (*n* = 386), *Pseudomonadota* (*n* = 242), *Bacteroidota* (*n* = 33), *Bacillota* A (*n* = 12), *Fusobacteriota* (*n* = 2), and *Patescibacteria* (*n* = 1). Additionally, we sequenced and assembled the genomes of 373 bacterial strains isolated from cheese, encompassing three phyla, namely *Pseudomonadota* (*n* = 195), *Actinomycetota* (*n* = 172) and *Bacillota* (*n* = 6). Using the GTDB classification, we found that 221 MAGs, encompassing 46 genera, as well as 44 bacterial strains, encompassing eight genera, could not be assigned to a species (<95% ANI) and thus represented potential new species (Figure 4). dRep clustering of these MAGs and genomes revealed potentially 81 new species. These species were particularly numerous among the genera *Halomonas* (23 MAGs and 11 isolates), *Psychrobacter* (23 MAGs and 6 isolates), *Brachybacterium* (10 MAGs and 14 isolates) and *Pseudoalteromonas* (10 MAGs and 3 isolates). Among the MAGs, 14 could not be assigned to the genus level. It was also the case for one *Actinomycetota* strain. We also compared our genomic dataset to a recent and large genomic database, namely curatedFoodMetagenomicData (cFMD) v1.1.0, which comprised 10,112 MAGs reconstructed from 2,533 food-associated metagenomes ^6^. Again, using an ANI threshold of 95% similarity ^62^, we found that 72 MAGs and 41 strains from our study remained potentially new and corresponded to 63 new species clusters (<95% ANI). They were particularly prevalent among *Pseudomonadota* (31 MAGs and 11 isolates) and *Actinomycetota* (28 MAGs and 29 isolates), confirming the high proportion of novel genomes in our dataset (Figure 5).

**Figure 4.**
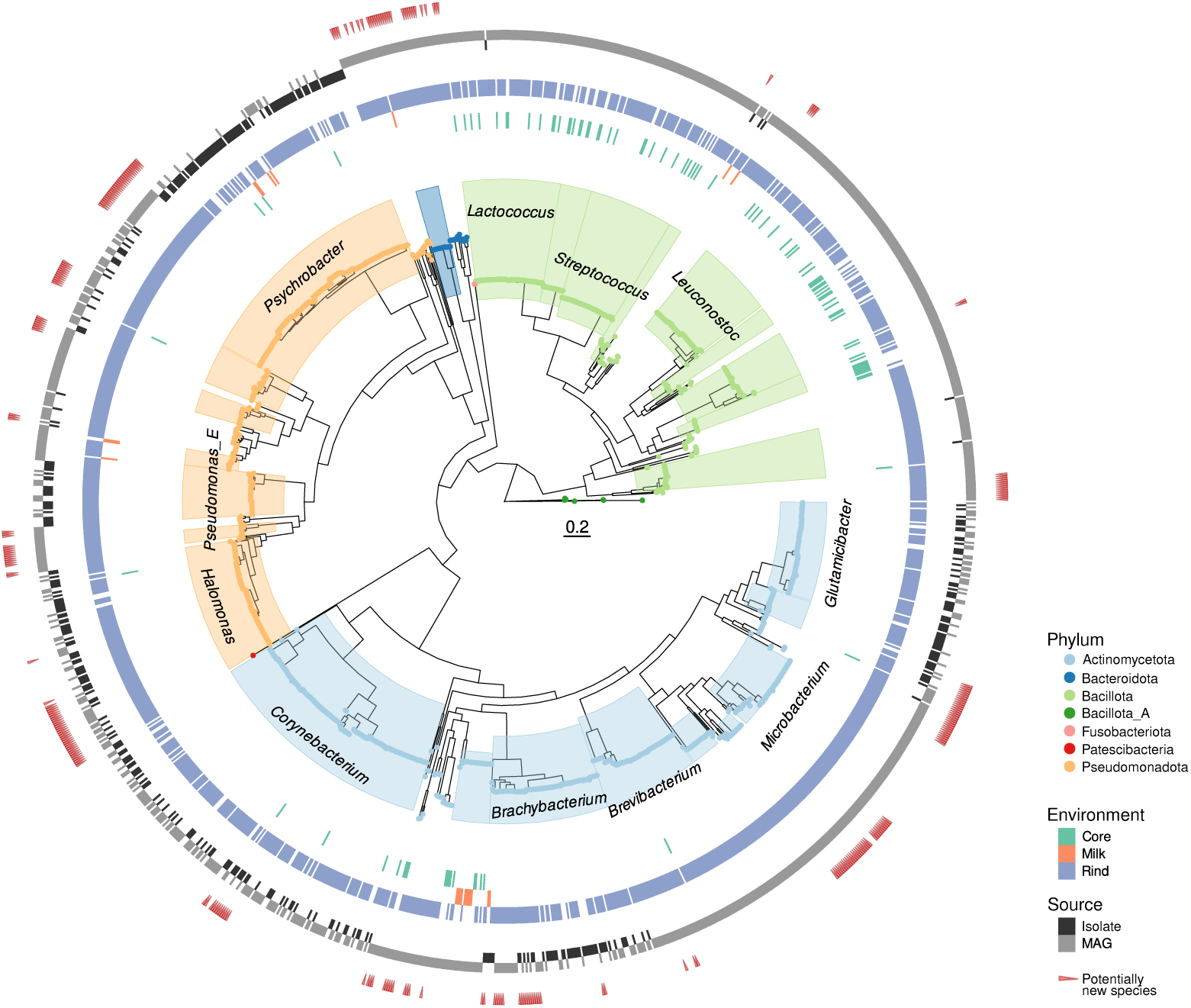
Bacterial genomic diversity. Maximum likelihood tree of the bacterial MAGs and isolate genomes generated in this study. Red triangles show MAGs/genomes not assigned to the species level by GTDB-tk.

**Figure 5.**
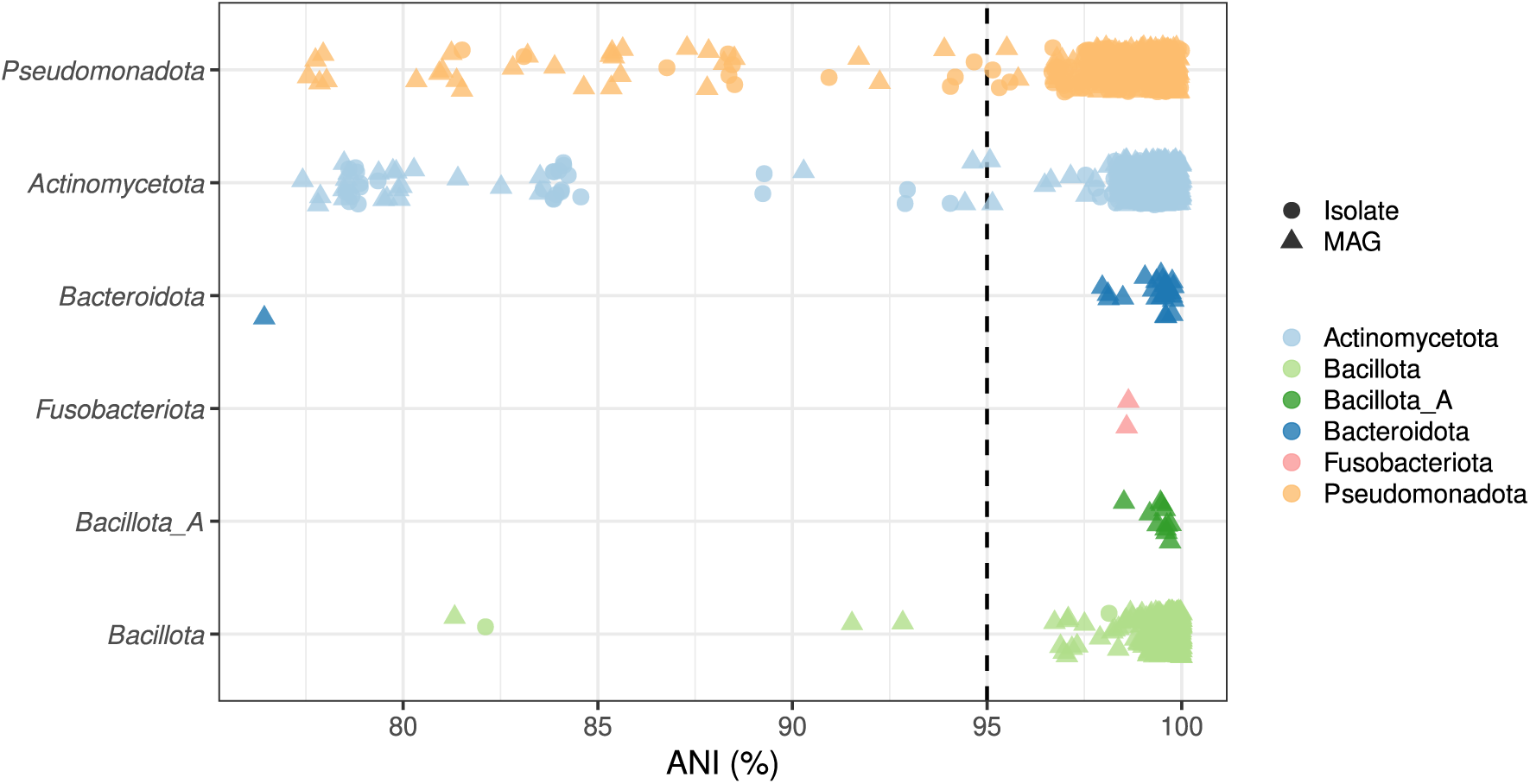
New bacterial genomic. Average nucleotide identity (ANI) between the bacterial MAGs and genomes of isolates generated in this study and the 10,112 MAGs of the cFMD database v1.1.0 ^6^. For 14 MAGs, no ANI value was reported by fastANI since it was below 80%. The vertical dotted line corresponds to the standard ANI cutoff (95%) used for species demarcation ^62^.

### Protein catalog

Then, we aggregated all our data to assemble a cheese protein catalog based on protein-level assembly of our metagenomes clustered (90% similarity) with the proteins predicted from our bacterial strains and MAGs (Figure 6). We obtained a total of 2.62 ✕ 10⁷ protein clusters (average length: 262 amino acids), including 4.76 ✕ 10⁶ clusters estimated to be complete proteins (18.15%). Rarefaction curves appeared to be unsaturated when considering either the whole catalog or bacterial or eukaryotic sequences, indicating that despite our sequencing effort there are still new proteins to discover in French PDO cheeses (Figure 6A). However, excluding singletons resulted in curves reaching a plateau and thus highlighting the role of unique proteins in the diversity of our catalog. A similar trend was observed when considering only sequences with at least 100 amino acids, indicating that the presence of short proteins did not bias our view of the catalog (Figure S4). Protein uniqueness was also clearly visible when looking at the protein cluster size distribution with a total of 14,926,596 singletons (56.87% of the protein clusters) (Figure 6B). Taxonomically, it was possible to assign 9,891,808 protein clusters to the bacterial domain (37.69%), 2,948,481 to eukaryotes (11.23%) and 132,700 to viruses (0.51%) (Figure 6C). Within this catalog, 13,232,482 protein clusters were also successfully assigned to a function (50.42%) using EggNOG (Figure 6D). These clusters encompassed cellular processes and signalling, information storage and processing, as well as metabolisms. In the latter one, transport and metabolism of amino acids, inorganic ions and carbohydrates were among the most diverse protein clusters. Noteworthy, among the annotated proteins, 1,935,053 protein clusters (14.62%) were assigned to existing but unknown functions.

**Figure 6.**
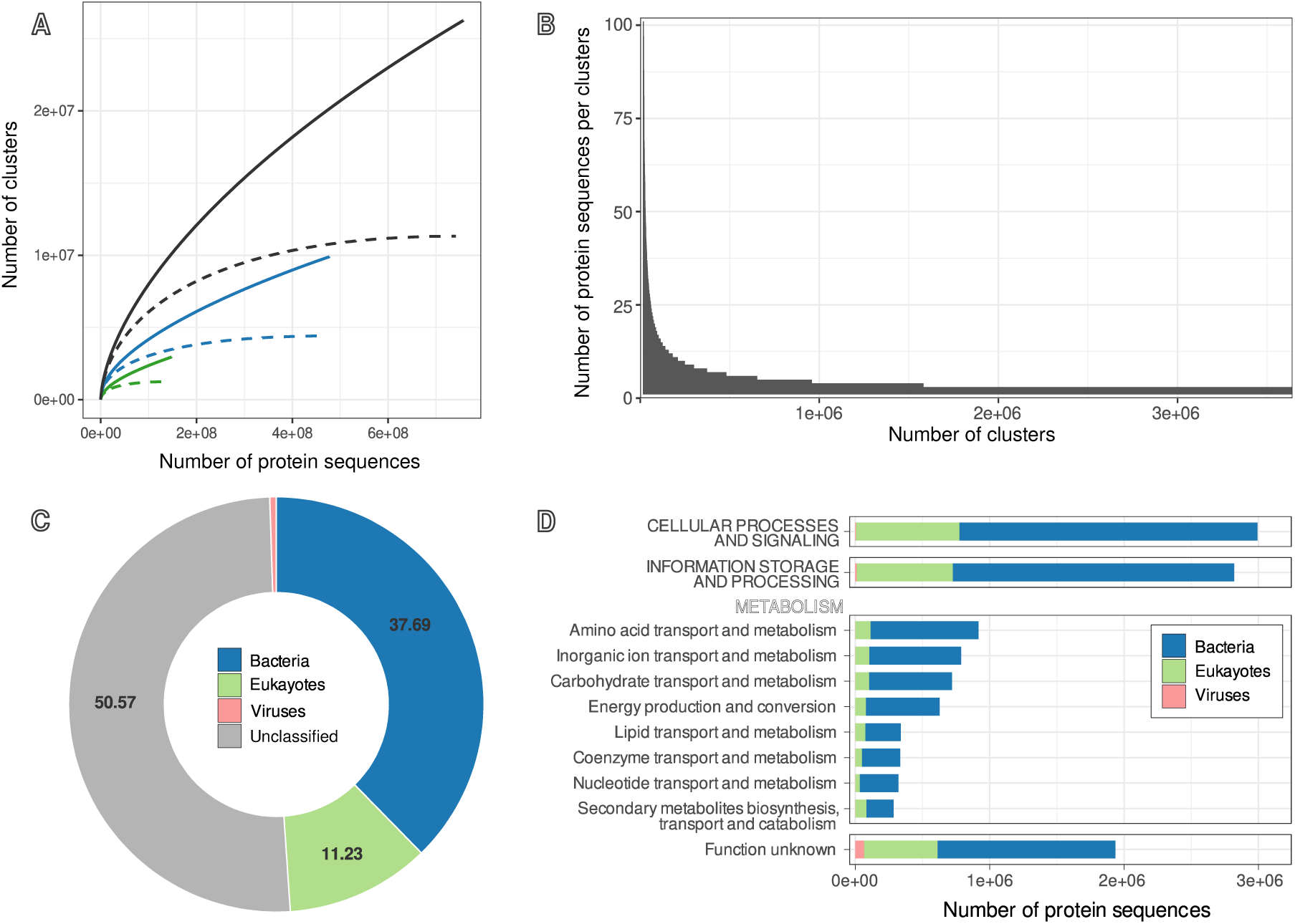
Protein catalog. **A.** Rarefactions curves (number of protein clusters as a function of the number of protein sequences). Black lines represent the total dataset, blue ones the bacterial proteins and green ones the eukaryotic proteins. Dashed lines represent curves without singletons. **B.** Histogram showing the distribution of the number of sequences within a 90% protein cluster. **C.** Taxonomic and **D.** functional distribution of the protein cluster catalog.

The catalog was also used to compare the three studied compartments (Table S4). Indeed, we found a set of 13 core protein clusters (10 bacterial proteins, one fungal, one bacteriophage protein and one unknown) common to our 146 samples and thus encompassing all milk, core and rind samples. When focusing on the 115 rind samples, this set increased to 87 core protein clusters, including 72 clusters of bacterial origin, 10 fungal, two bacteriophage and three unclassified protein clusters. Besides general cellular processes, we also found proteins involved in various metabolisms such as carbohydrates (*e.g.* glycosyl transferase and hydrolase), nucleic acids (*e.g.* DNA gyrase and nuclease) and amino acids (*e.g.* endopeptidase lactocepin and peptide permease).

### The case of methionine gamma-lyase

To demonstrate the potential of our dataset, we investigated the distribution and diversity of methionine gamma-lyase (MGL, also referred as L-methioninase, EC 4.4.1.11), an enzyme involved in the degradation of methionine into methanethiol, a volatile sulfur compound that can further be oxidized to dimethyldisulfide (DMDS) and dimethyltrisulfide (DMTS). These three compounds play a significant role in cheese flavor ^74^. We found 50 protein clusters corresponding to 1,899 sequences (1,840 in the surface, 28 in the core, 31 in the milk) in our protein catalog, as well as 41 MGL sequences from MAGs and 9 from bacterial isolate genomes (Figure S5). Each genome possesses one copy of MGL, except for the one assigned to *Fusobacterium,* which possesses two copies. The distribution and prevalence of this enzyme on cheese rind greatly differ between technologies, with MGL being present in all lactic washed rind cheeses and highly abundant in various soft washed rind cheeses (Figure 7A). On the other hand, MGL was only detected in one sample of hard-cooked cheese and two internal blue mold cheeses. Taxa harboring MGL were restricted to three phyla, namely *Pseudomonadata* (95.42% of the MGL sequences)*, Fusobacteria* (2.37%) and *Bacillota* (2.21%). In particular, we found *Pseudoalteromonas* (*n*=729 sequences), *Serratia* (*n*=444)*, Pseudomonas* (*n*=254)*, Hafnia* (*n*=159) and *Proteus* (*n*=124) among the most important genera carrying MGL (Figure 7B).

**Figure 7.**
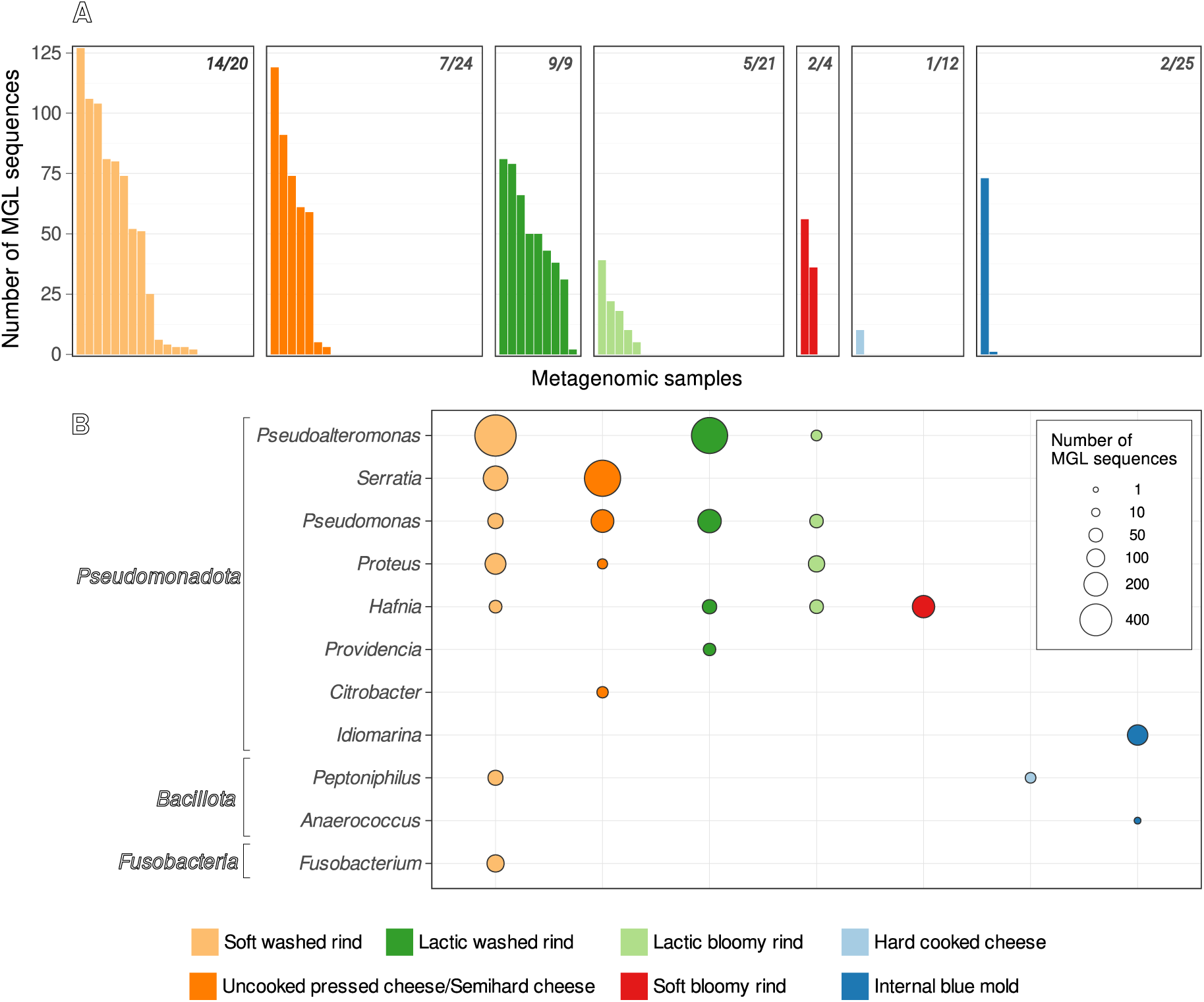
Diversity of methionine gamma-lyase (MGL) in the different cheese technologies recovered from our protein catalog. **A.** Distribution of the number of MGL sequences in the cheese rind samples. One bar represents the number of MGL sequences in a metagenomic sample. Ratio on the left top of each plot corresponds to the prevalence of MGL in each cheese technology. **B.** Taxonomic distribution of the MGL in the different cheese rinds.

## Discussion

Combining culturomics and metagenomics, our study provides a wide (meta)genomic resource of cheese-associated microorganisms encompassing a large diversity of French PDO-ripened cheeses. This national-scale survey reveals distinct ecological patterns, an overview of the fungal, bacterial and viral diversity present in French cheeses, potentially new bacterial species that will be formally described in due course, and a huge set of functions to be further explored.

### Ecological patterns in French PDO cheeses

After ensuring that our dairy samples were sufficiently sequenced to properly cover the sequence diversity (Figure S1A), we used the Nd metric to compare sequence diversity across samples. Sequence diversity was higher in rind than in core and milk samples (Figure S1B), highlighting the taxonomic and functional complexity of the cheese ripening microbiome ^75,76^ and the necessity to further explore the rind habitat. We also found a positive correlation between cheese pH and sequence diversity (Figure S2), a trend that was already reported in previous work on cheese microbiome ^21^.

Then, we investigated more precisely the ecological patterns underlying the diversity of bacteria, eukaryotes, viruses but also microbial functions. For these four categories, we consistently observed a significant effect of the environment (milk, core, rind) on both the *alpha* and *beta* diversity (Figures 2, 3 and S1). We observed a higher taxonomic richness in rind than in core for many samples, as previously reported in the literature ^77^. Surprisingly, we found a lower diversity in milk compared to cheese samples, which contradicts metabarcoding results from the same samples ^22^. This could be explained by the absence of PCR amplification and the lower microbial biomass in milk ^78^ which could have prevented the detection of rare taxa. Our study also revealed significant differences in functional diversity between sample types (milk, core, rind) but also between cheese-making technologies. A previous study ^79^ also reported significant functional differences between core and rind but failed to detect the cheese-making technology effect.

Indeed, our analysis showed that the ch*e*ese technological family significantly influenced both *alpha* and *beta* diversity in the rind, confirming results observed by a metabarcoding approach targeting bacterial and fungal communities on similar French PDO cheese samples ^22^, as well as other studies performed on different cheeses ^19^. Our work also extends on this technological effect for viral (Figure 2) and functional diversity (Figure 3). Interestingly, we found that functional genes (Figure 3B) explained more variance than taxonomic marker genes (Figures 2B, E, H) for both environmental and technological effects. A similar finding was also reported for marine metagenomics ^80^, suggesting that functional genes can be better predictors of microbial assembly than taxonomic ones.

Among the discriminant metabolic functions between cheese rind technologies, we identified metabolic categories of interest for cheese makers (Figure 3D), like proteolysis and lipolysis ^81,82^. Indeed, the amino acid transport and metabolism category was driven mainly by variations in the abundance of genes involved in oligopeptide import (COG4166) which could allow microorganisms to metabolize the products of proteolysis as well as in cysteine/methionine metabolism and/or biosynthesis of secondary metabolites (COG0527, COG0345, COG1760, COG0133, COG2423, COG1168) which could participate to cheese flavor production. Regarding the lipid category, the two most abundant genes discriminating between cheese technologies encoded two acyl-CoA thioesterases (COG1946, COG1607), which are enzymes involved in the hydrolysis of acyl-coenzyme A to the free fatty acid and coenzyme A, and thus in lipolysis ^83^.

### A metagenome-based taxonomic survey

Using various genomic markers, we survey the taxonomy and abundance of bacteria, eukaryotes and viruses, without the inherent biases of the metabarcoding approach ^84^. Our analysis confirms the absence of archaea in cheese, as well as the high taxonomic microbial diversity in cheese rind compared to cheese core. Among bacteria, lactic-acid bacteria (LAB) were among the most prevalent and abundant species in the dataset (seven LAB species with more than 1% abundance), with *Streptococcus thermophilus* and *Lactococcus lactis* being the two most abundant species. The LAB were also quite diverse with 29 species of *Lactobacillaceae*, 18 of *Streptococcaceae* and 8 of *Leuconostocaceae* encompassing mesophilic and thermophilic starters, as well as non-starter LAB containing mesophilic obligate and facultative hetero-fermentative lactobacilli ^85^. Ripening bacteria were even more diverse with 102 species of *Gammaproteobacteria* and 50 species of *Actinomycetia.* Among them, *Corynebacterium variabile, Corynebacterium casei, Brevibacterium linens, Microbacterium gubbeenense, Glutamicibacter arilaitensis (Actinomycetia), Hafnia alvei and Psychrobacter* sp*. (Gammaproteobacteria)* were among the most abundant species.

Among eukaryotes, we identified well-known cheese fungi with high prevalence and abundance (>3% of the total eukaryotic abundance) such as *Debaryomyces hansenii, Diutina catenulata, Kluyveromyces lactis, Penicillium roqueforti* and *Candida sake* ^86^. We also identified various low abundance and low prevalence fungi with environmental origins, such as three species of *Fusarium* (potential plant pathogens), one of *Cordyceps* (arthropod pathogen) and various yeasts not described as inoculated by cheese-makers (*Saturnispora silvae, Teunomyces cretensis, Meyerozyma caribbica, Saturnispora mendoncae*). *Saturnispora mendoncae* was already isolated from artisanal French cheese ^87^. However, the role of these non-inoculated yeasts in the cheese ecosystem remains to be elucidated. Interestingly, our metagenomic survey also detected the presence of the mite *Tyrophagus putrescentiae*, an acarid frequently found on cheese ^88,89^.

We also recovered viral sequences matching those of phages infecting ripening bacteria already isolated from the rind of a French smear-ripened cheese (Paillet *et al.*, 2022) and a canadian washed-rind cheese ^91^. Among those phages, *Leuconostoc* phage Diderot was the most widespread in French PDO cheeses, detected in 27 samples from 16 distinct PDO cheeses. *Brevibacterium* phage Rousseau was detected in 10 samples from 4 PDO cheeses, and *Glutamicibacter* phage Montesquieu in 4 samples from 3 PDO cheeses. Both *Glutamicibacter* phage Voltaire and *Psychrobacter* phage d’Alembert were exclusively detected in a single PDO cheese technology. Our findings highlight the widespread distribution of such phages in traditional cheeses and point to the potential discovery of hundreds of novel ones, since several vOTUs detected in our metagenomic survey are currently classified as unknown *Caudoviricetes* and do not match the genomes of common dairy phages.

### A bacterial genome catalog from French PDO cheeses

Combining metagenomics and culturomics, we generated 1,119 bacterial MAGs and 373 genomes of bacterial strains from cheese (Figure 4) and that covered most of the prokaryotic diversity described in French PDO cheeses, namely members of *Actinomycetota, Bacillota* and *Pseudomonadota* ^22^. In particular, the use of a culture medium supplemented with NaCl allowed the isolation of various halophilic and halotolerant bacteria that are known to be prevalent in cheese ^92^, for instance members of the halotolerant genus *Psychrobacter* (*n* = 118 isolates) ^93^, and of the halophilic or halotolerant genus *Halomonas* (*n* = 57 isolates) ^94^. The diversity of these genera was also recovered with the MAGs (60 *Psychrobacter* and 43 *Halomonas*), confirming their importance. These bacterial genomes are valuable resources for further pangenome analysis, especially to understand the adaptation to the cheese habitat as well as the metabolic potential of cheese ripening bacteria ^95,96^. Additionally these genomes will facilitate sub-species analyses ^27^.

Our study also identified some genomic novelty in the cheese ecosystem. Indeed, compared to GTDB ^46^ or to the food specific database cFMD ^6^, many taxa found in both MAGs and genomes of isolates represent potentially new species, namely 81 potentially new species when compared to GTDB. Another metagenomic study of cheese samples collected across multiple countries and continents also highlighted the presence of previously uncharacterized species in cheese, albeit to a lesser extent, identifying 22 potentially novel species out of 328 MAGs reconstructed from 184 samples ^79^. This clearly demonstrates that traditional cheeses harbor a high genomic diversity with bacterial species yet to be described, and thus contributing, as reservoirs, to the hidden diversity in food ^27,28^. These genomes might also conceal a biotechnological potential and as such, the strains would be a valuable resource for local cheese producers ^32,33^ but also to better understand the typicity of PDO cheeses.

Among our isolates, we also obtained strains previously uncultured (no type material nor bacterial isolate available to date) which are essential for the phenotypic characterisation of these organisms and which would be precious for the formal description of new bacterial taxa. This was for instance the case of *Corynebacterium faecigallinarum*, which was found in chicken gut metagenomes but not isolated so far ^97^.

On a side note, we found in our dataset 28 MAGs assigned to *Cellulosimicrobium funkei*. Their unexpected presence in food samples, combined with their occurrence in multiple and distinct samples, suggests they are a potential contaminant of the DNA extraction procedure, more precisely from the enzyme mix of lyticase (Sigma-Aldrich L4025). Therefore, these genomes should be used in further studies as reference genomes to filter reads during the bioinformatic procedure of metagenomic samples. On the other hand, MAG analysis also unveiled the identification of potential biotic associations in cheese. Indeed, we recovered two MAGs from the same cheese rind sample (AOP1_E1_surf): a MAG belonging to *Saccharimonadales* UBA10027 (*Patescibacteria*) and a MAGassigned to *Pauljensenia* sp (*Actinobacteriota)*. Interestingly, members of *Saccharimonadia* (formerly classified as TM7) have been shown to be an epibiont of *Pauljensenia odontolyticus* (formerly known as *Actinomyces odontolyticus*) in the human oral cavity ^98^. Future work should include microscopy to validate the existence of a similar association in cheese. If validated, it could add a new hypothesis about the habitat transition in the *Saccharibacteria* ^99^, with cheese serving as an intermediate between the environment and the human oral cavity. Noteworthy, the 16S rRNA gene metabarcoding approach failed to detect any *Saccharimonadia* reads from the same cheese samples^22^, highlighting the benefits of shotgun metagenomics.

### A protein catalog to explore cheese functions

By combining protein sequences assembled directly from metagenomes with protein sequences predicted from genomes, we built a taxonomically and functionally annotated protein catalog composed of 26.2 millions protein clusters. As such, this catalog will give the opportunity to address with confidence the taxonomic origin of many proteins. Indeed, the inclusion of the genome-level data from isolates and MAGs significantly increases the taxonomic resolution of the annotation. This catalog will also be useful for protein-centric analyses ^100^.

The distribution of the number of sequences per protein cluster and the rarefaction curves of our catalog (Figure 6) revealed similar patterns to those found in catalogs from other ecosystems. Indeed, as shown in a global survey, most gene-coding proteins are rare^1^. In the case of French PDO cheeses, these rare proteins could partly belong to low abundant species that can play key roles in such ecosystems ^22^. Therefore, careful exploration of these proteins could be relevant to better understand the cheese ecosystem’s functioning, especially considering functional rarity ^101^.

Although no core microbiota for bacteria and fungi was detected in PDO cheese using a metabarcoding approach ^22^, we identified here a set of 13 core protein clusters encompassing viral, bacterial and fungal origins. It suggests a functional continuum from milk to cheese. Focusing only on the rind, this core extends to 87 core protein clusters, potentially reflecting the common properties found in cheese rind habitat (pH, presence of oxygen, sources of carbon and nitrogen).

Taxonomic and functional annotation of our cheese protein catalog revealed that around half of the protein clusters are taxonomically and functionally unknown. Although some sequences could be too short to be assigned with confidence, many functions remain to be discovered in the cheese microbiome. This could be achieved by either *in vitro* characterisation of the unknown proteins or by improving the protein annotation ^102^. The idea of unexplored diversity is also supported by the unsaturated rarefaction curves, suggesting that sampling more metagenomes will bring more new rare proteins (Figure 6).

As illustrated by the example of methionine gamma-lyase (Figure 7), our catalog can be easily used to explore functional diversity in cheese and to identify microbial taxa involved in specific pathways ^103^. The cheese ecosystem is known to harbor MGL producers ^104^. Our survey allowed us to clarify the taxonomy and abundance of MGL-producing bacteria. Among the genera of MGL producers detected in our catalog, some were already described in the literature ^105^ such as *Serratia* ^106^, *Pseudomonas* ^107^, *Hafnia* ^108^, *Citrobacter* ^109^, *Peptoniphilus* ^110^ and *Fusobacterium* ^111^. However, to the best of our knowledge, *Pseudoalteromonas* has not yet been reported as a MGL producer and might represent a new player in sulfur metabolism on cheese rind. *Pseudoalteromonas* is known to be present in various cheeses ^112^ but its role in cheese ripening remains obscure. This example shows how this catalog can be used to improve our knowledge on the role of cheese ripening taxa.

More generally, our catalog will be a valuable resource for data mining. Examples of applications could be in the emerging field of synthetic ecology. Indeed, searching for functions in the annotated genomes could be a first step to design synthetic microbial communities (SynComs) in order to improve the functional properties of fermented foods ^113–115^. Another example, our catalog could also be used for taxonomical and functional annotation of metaproteomic datasets ^116^.

## Supporting information

Suuplementary Figure 1

Suuplementary Figure 2

Suuplementary Figure 3

Suuplementary Figure 4

Suuplementary Figure 5

## Acknowledgments

We are grateful to the INRAE MIGALE bioinformatics facility (MIGALE, INRAE, 2020. Migale bioinformatics Facility, doi: 10.15454/1.5572390655343293E12) for providing help, computing and storage resources. We also thank Anne-Sophie Sarthou, Lucie Arnoult and Margot Michottey for the technical assistance.

## Data availability

Raw metagenomic data are available under the Bioproject PRJEB64630.

Raw bacterial genomic data are available under the Bioproject PRJEB64629.

Assembled bacterial genomes are available under the Bioproject PRJNA1308522.

Bacterial MAGs are available under the Bioproject PRJEB64630.

The protein catalog and the scripts are available at: https://entrepot.recherche.data.gouv.fr/dataverse/metapdocheese_metagenomics_culturomics/

## Additional Files

**Figure S1.** Estimated coverage (A) and sequence diversity Nd (B) of the 146 metagenomic samples, based on Nonpareil v3.4.1. Statistical comparisons were computed with Kruskal-Wallis test followed by Dunn’s test with Bonferroni adjustment.

**Figure S2.** Relationship between the sequence diversity Nd and cheese pH for (A) all the cheese samples, (B) only the rind samples and (C) only the core samples. The blue line represents the fitted linear equation. Pearson correlation coefficients (R) are presented for each plot.

**Figure S3.** Barplots showing the average taxonomic distribution at the contig level within the samples. Classification was done using MMseqs2.

**Figure S4.** Effect of protein length and singletons on the rarefaction curves of the protein catalog. The light grey line represents all the protein clusters while the dark grey line excludes the singletons. The orange line represents the protein clusters with a length of at least 100 amino acids while the dark red line represents the protein clusters with a length of at least 100 amino acids and without singleton.

**Figure S5.** Maximum likelihood phylogenetic tree of methionine-gamma lyase sequences. A total of 1251 amino-acid reference sequences from InterPro (prefix reference_), 50 sequences from bacterial MAG or genome strains (prefix genome_) and 50 sequences from PLASS (prefix plass_) were aligned with MAFFT. The tree was inferred with IQ-TREE2 using the LG+I+G4 model of amino-acid evolution (based on ModelFinder selection).

**Table S1.** Summary of the 146 metagenome characteristics.

**Table S2.** Summary of the mOTUS, EukDetect and vOTUs results.

**Table S3.** Summary of the MAG and bacterial strain genome properties.

**Table S4.** List of the protein clusters shared between all samples or only between the cheese rind samples.

## References

1. Coelho, L. P. et al. Towards the biogeography of prokaryotic genes. Nature 601, 252–256 (2022).

2. Almeida, A. et al. A unified catalog of 204,938 reference genomes from the human gut microbiome. Nature Biotechnology 39, 105–114 (2021).

3. Nayfach, S. et al. A genomic catalog of Earth’s microbiomes. Nature Biotechnology 39, 499–509 (2021).

4. Garner, R. E. et al. A genome catalogue of lake bacterial diversity and its drivers at continental scale. Nat Microbiol 1–15 (2023) doi:10.1038/s41564-023-01435-6.

5. Wu, D., Seshadri, R., Kyrpides, N. C. & Ivanova, N. N. A metagenomic perspective on the microbial prokaryotic genome census. Science Advances 11, eadq2166 (2025).

6. Carlino, N. et al. Unexplored microbial diversity from 2,500 food metagenomes and links with the human microbiome. Cell 187, 5775–5795.e15 (2024).

7. Dunn, R. R., Wilson, J., Nichols, L. M. & Gavin, M. C. Toward a global ecology of fermented foods. Current Anthropology 62, S220–S232 (2021).

8. Miller, V. et al. Global, regional, and national consumption of animal-source foods between 1990 and 2018: findings from the Global Dietary Database. The Lancet Planetary Health 6, e243–e256 (2022).

9. Button, J. E. & Dutton, R. J. Cheese microbes. Current Biology 22, R587–R589 (2012).

10. Donnelly, C. W. From Pasteur to probiotics: A historical overview of cheese and microbes. Microbiology Spectrum 1, 10.1128/microbiolspec.cm-0001–0012 (2013).

11. Almena-Aliste, M. & Mietton, B. Cheese classification, characterization, and categorization: A global perspective. Microbiology Spectrum 2, 10.1128/microbiolspec.cm-0003–2012 (2014).

12. Montel, M.-C. et al. Traditional cheeses: Rich and diverse microbiota with associated benefits. International Journal of Food Microbiology 177, 136–154 (2014).

13. Fox, P. F. & McSweeney, P. L. H. Chapter 1 – Cheese: An Overview. in Cheese (Fifth Edition) (eds McSweeney, P. L. H., Cotter, P. D., Everett, D. W. & Govindasamy-Lucey, Rani) 5–20 (Academic Press, San Diego, 2025). doi:10.1016/B978-0-12-417012-4.00001-6.

14. McSweeney, P. L. H. & Fox, P. F. Chapter 30 – Diversity and classification of cheese varieties: An Overview. in Cheese (Fifth Edition) (eds McSweeney, P. L. H., Cotter, P. D., Everett, D. W. & Govindasamy-Lucey, Rani) 771–797 (Academic Press, San Diego, 2025). doi:10.1016/B978-0-12-417012-4.00031-4.

15. Ropars, J., Cruaud, C., Lacoste, S. & Dupont, J. A taxonomic and ecological overview of cheese fungi. International Journal of Food Microbiology 155, 199–210 (2012).

16. Paillet, T., Lamy-Besnier, Q., Figueroa, C., Petit, M.-A. & Dugat-Bony, E. Dynamics of the viral community on the surface of a French smear-ripened cheese during maturation and persistence across production years. mSystems 0, e00201–24 (2024).

17. Brennan, N. M. et al. Biodiversity of the bacterial flora on the surface of a smear cheese. Applied and Environmental Microbiology 68, 820–830 (2002).

18. Irlinger, F., Layec, S., Hélinck, S. & Dugat-Bony, E. Cheese rind microbial communities: diversity, composition and origin. FEMS Microbiology Letters 362, 1–11 (2015).

19. Wolfe, B. E., Button, J. E., Santarelli, M. & Dutton, R. J. Cheese rind communities provide tractable systems for in situ and in vitro studies of microbial diversity. Cell 158, 422–433 (2014).

20. Raymond-Fleury, A. et al. Analysis of microbiota persistence in Quebec’s terroir cheese using a metabarcoding approach. Microorganisms 10, 1381 (2022).

21. Reuben, R. C., Langer, D., Eisenhauer, N. & Jurburg, S. D. Universal drivers of cheese microbiomes. iScience 26, 105744 (2023).

22. Irlinger, F. et al. A comprehensive, large-scale analysis of “terroir” cheese and milk microbiota reveals profiles strongly shaped by both geographical and human factors. ISME Communications 4, ycae095 (2024).

23. Louw, N. L., Lele, K., Ye, R., Edwards, C. B. & Wolfe, B. E. Microbiome assembly in fermented foods. Annual Review of Microbiology 77, 381–402 (2023).

24. Kastman, E. K. et al. Biotic Interactions Shape the Ecological Distributions of Staphylococcus Species. mBio 7, e01157–16 (2016).

25. Cosetta, C. M., Kfoury, N., Robbat, A. & Wolfe, B. E. Fungal volatiles mediate cheese rind microbiome assembly. Environmental Microbiology 1462-2920.15223 (2020) doi:10.1111/1462-2920.15223.

26. Mekuli, R. et al. Iron-based microbial interactions: the role of iron metabolism in the cheese ecosystem. Journal of Bacteriology 207, e00539–24 (2025).

27. Flörl, L., Meyer, A. & Bokulich, N. A. Exploring sub-species variation in food microbiomes: a roadmap to reveal hidden diversity and functional potential. Applied and Environmental Microbiology 91, e00524–25 (2025).

28. Pasolli, E. & Ercolini, D. The hidden diversity of food microbiomes. Current Opinion in Food Science 64, 101318 (2025).

29. Fontana, F. et al. Multifactorial microvariability of the Italian raw milk cheese microbiota and implication for current regulatory scheme. mSystems 8, e01068–22 (2023).

30. Salamandane, A. et al. Metagenomic analysis of the bacterial microbiome, resistome and virulome distinguishes Portuguese Serra da Estrela PDO cheeses from similar non-PDO cheeses: An exploratory approach. Food Research International 189, 114556 (2024).

31. Santamarina-García, G. et al. Shotgun metagenomic sequencing reveals the influence of artisanal dairy environments on the microbiomes, quality, and safety of Idiazabal, a raw ewe milk PDO cheese. Microbiome 12, 262 (2024).

32. Grizon, A. et al. Genetic and technological diversity of Streptococcus thermophilus isolated from the Saint-Nectaire PDO cheese-producing area. Frontiers in Microbiology 14, (2023).

33. Grizon, A. et al. Genomic characterization of wild *Lactobacillus delbrueckii* strains reveals low diversity but strong typicity. Microorganisms 12, 512 (2024).

34. Alberti, A. et al. Viral to metazoan marine plankton nucleotide sequences from the Tara Oceans expedition. Sci Data 4, 170093 (2017).

35. Li, R., Li, Y., Kristiansen, K. & Wang, J. SOAP: short oligonucleotide alignment program. Bioinformatics 24, 713–714 (2008).

36. Chen, S., Zhou, Y., Chen, Y. & Gu, J. fastp: an ultra-fast all-in-one FASTQ preprocessor. Bioinformatics 34, i884–i890 (2018).

37. Wick, R. R. et al. Unicycler: Resolving bacterial genome assemblies from short and long sequencing reads. PLOS Computational Biology 13, e1005595 (2017).

38. Gurevich, A., Saveliev, V., Vyahhi, N. & Tesler, G. QUAST: quality assessment tool for genome assemblies. Bioinformatics 29, 1072–5 (2013).

39. Li, H. & Durbin, R. Fast and accurate short read alignment with Burrows-Wheeler transform. Bioinformatics 25, 1754–60 (2009).

40. Nurk, S., Meleshko, D., Korobeynikov, A. & Pevzner, P. A. metaSPAdes: a new versatile metagenomic assembler. Genome Research 27, 824–834 (2017).

41. Mikheenko, A., Saveliev, V. & Gurevich, A. MetaQUAST: evaluation of metagenome assemblies. Bioinformatics 32, 1088–1090 (2016).

42. Tadrent, N., Dedeine, F. & Hervé, V. SnakeMAGs: a simple, efficient, flexible and scalable workflow to reconstruct prokaryotic genomes from metagenomes. F1000Res 11, 1522 (2023).

43. Kang, D. D. et al. MetaBAT 2: an adaptive binning algorithm for robust and efficient genome reconstruction from metagenome assemblies. PeerJ 7, e7359 (2019).

44. Orakov, A. et al. GUNC: detection of chimerism and contamination in prokaryotic genomes. Genome Biology 22, 178 (2021).

45. Chaumeil, P.-A., Mussig, A. J., Hugenholtz, P. & Parks, D. H. GTDB-Tk v2: memory friendly classification with the Genome Taxonomy Database. 2022.07.11.499641 Preprint at 10.1101/2022.07.11.499641 (2022).

46. Parks, D. H. et al. GTDB: an ongoing census of bacterial and archaeal diversity through a phylogenetically consistent, rank normalized and complete genome-based taxonomy. Nucleic Acids Research 50, D785–D794 (2022).

47. Olm, M. R., Brown, C. T., Brooks, B. & Banfield, J. F. dRep: a tool for fast and accurate genomic comparisons that enables improved genome recovery from metagenomes through de-replication. The ISME Journal 11, 2864–2868 (2017).

48. Seemann, T. Prokka: rapid prokaryotic genome annotation. Bioinformatics (Oxford, England) 30, 2068–9 (2014).

49. Cantalapiedra, C. P., Hernández-Plaza, A., Letunic, I., Bork, P. & Huerta-Cepas, J. eggNOG-mapper v2: Functional annotation, orthology assignments, and domain prediction at the metagenomic scale. Molecular Biology and Evolution 10.1093/MOLBEV/MSAB293 (2021) doi:10.1093/MOLBEV/MSAB293.

50. Huerta-Cepas, J. et al. Fast genome-wide functional annotation through orthology assignment by eggNOG-mapper. Molecular Biology and Evolution 34, (2017).

51. Ruscheweyh, H.-J. et al. Cultivation-independent genomes greatly expand taxonomic-profiling capabilities of mOTUs across various environments. Microbiome 10, 212 (2022).

52. Lind, A. L. & Pollard, K. S. Accurate and sensitive detection of microbial eukaryotes from whole metagenome shotgun sequencing. Microbiome 9, 58 (2021).

53. Kieft, K., Zhou, Z. & Anantharaman, K. VIBRANT: automated recovery, annotation and curation of microbial viruses, and evaluation of viral community function from genomic sequences. Microbiome 8, 90 (2020).

54. Nayfach, S. et al. CheckV assesses the quality and completeness of metagenome-assembled viral genomes. Nat Biotechnol 39, 578–585 (2021).

55. Roux, S. et al. Minimum Information about an Uncultivated Virus Genome (MIUViG). Nat Biotechnol 37, 29–37 (2019).

56. Steinegger, M. & Söding, J. MMseqs2 enables sensitive protein sequence searching for the analysis of massive data sets. Nat Biotechnol 35, 1026–1028 (2017).

57. Camargo, A. P. et al. Identification of mobile genetic elements with geNomad. Nat Biotechnol 42, 1303–1312 (2024).

58. Camacho, C. et al. BLAST+: architecture and applications. BMC Bioinformatics 10, 421 (2009).

59. Asnicar, F., Weingart, G., Tickle, T. L., Huttenhower, C. & Segata, N. Compact graphical representation of phylogenetic data and metadata with GraPhlAn. PeerJ 3, e1029 (2015).

60. Minh, B. Q. et al. IQ-TREE 2: New models and efficient methods for phylogenetic inference in the genomic era. Molecular Biology and Evolution 37, 1530– 1534 (2020).

61. Yu, G., Smith, D. K., Zhu, H., Guan, Y. & Lam, T. T.-Y. <scp>ggtree</scp>: an <scp>r</scp> package for visualization and annotation of phylogenetic trees with their covariates and other associated data. Methods in Ecology and Evolution 8, 28–36 (2017).

62. Jain, C., Rodriguez-R, L. M., Phillippy, A. M., Konstantinidis, K. T. & Aluru, S. High throughput ANI analysis of 90K prokaryotic genomes reveals clear species boundaries. Nature Communications 9, 5114 (2018).

63. Steinegger, M., Mirdita, M. & Söding, J. Protein-level assembly increases protein sequence recovery from metagenomic samples manyfold. Nature Methods 16, 603–606 (2019).

64. Steinegger, M. & Söding, J. Clustering huge protein sequence sets in linear time. Nature Communications 9, 2542 (2018).

65. Schloss, P. D. et al. Introducing mothur: open-source, platform-independent, community-supported software for describing and comparing microbial communities. Applied and Environmental Microbiology 75, 7537–7541 (2009).

66. Suzek, B. E. et al. UniRef clusters: a comprehensive and scalable alternative for improving sequence similarity searches. Bioinformatics 31, 926–932 (2015).

67. Eddy, S. R. Accelerated profile HMM searches. PLoS Computational Biology 7, e1002195 (2011).

68. Paysan-Lafosse, T. et al. InterPro in 2022. Nucleic Acids Research 51, D418–D427 (2023).

69. Katoh, K. & Standley, D. M. MAFFT multiple sequence alignment software version 7: improvements in performance and usability. Molecular Biology and Evolution 30, 772–80 (2013).

70. Rodriguez-R, L. M., Gunturu, S., Tiedje, J. M., Cole, J. R. & Konstantinidis, K. T. Nonpareil 3: Fast estimation of metagenomic coverage and sequence diversity. mSystems 3, e00039–18 (2018).

71. Wickham, H. et al. Welcome to the Tidyverse. Journal of Open Source Software 4, 1686 (2019).

72. Fernandes, A. D. et al. Unifying the analysis of high-throughput sequencing datasets: characterizing RNA-seq, 16S rRNA gene sequencing and selective growth experiments by compositional data analysis. Microbiome 2, 15 (2014).

73. Gu, Z. Complex heatmap visualization. iMeta 1, e43 (2022).

74. Yvon, M. & Rijnen, L. Cheese flavour formation by amino acid catabolism. International Dairy Journal 11, 185–201 (2001).

75. Beresford, T. & Williams, A. The microbiology of cheese ripening. in Cheese: Chemistry, Physics and Microbiology (eds Fox, P. F., McSweeney, P. L. H., Cogan, T. M. & Guinee, T. P.) vol. 1 287–317 (Academic Press, 2004).

76. McSweeney, P. L. H. Biochemistry of cheese ripening. International Journal of Dairy Technology 57, 127–144 (2004).

77. Dugat-Bony, E. et al. Highlighting the microbial diversity of 12 French cheese varieties. International Journal of Food Microbiology 238, 265–273 (2016).

78. Franciosi, E., Settanni, L., Cologna, N., Cavazza, A. & Poznanski, E. Microbial analysis of raw cows’ milk used for cheese-making: influence of storage treatments on microbial composition and other technological traits. World J Microbiol Biotechnol 27, 171–180 (2011).

79. Walsh, A. M., Macori, G., Kilcawley, K. N. & Cotter, P. D. Meta-analysis of cheese microbiomes highlights contributions to multiple aspects of quality. Nat Food 1, 500–510 (2020).

80. Terzin, M. et al. Gene content of seawater microbes is a strong predictor of water chemistry across the Great Barrier Reef. Microbiome 13, 11 (2025).

81. Sousa, M. J., Ardö, Y. & McSweeney, P. L. H. Advances in the study of proteolysis during cheese ripening. International Dairy Journal 11, 327–345 (2001).

82. Collins, Y. F., McSweeney, P. L. H. & Wilkinson, M. G. Lipolysis and free fatty acid catabolism in cheese: a review of current knowledge. International Dairy Journal 13, 841–866 (2003).

83. Hunt, M. C. & Alexson, S. E. H. The role Acyl-CoA thioesterases play in mediating intracellular lipid metabolism. Progress in Lipid Research 41, 99–130 (2002).

84. Srinivas, M., O’Sullivan, O., Cotter, P. D., Sinderen, D. van & Kenny, J. G. The application of metagenomics to study microbial communities and develop desirable traits in fermented foods. Foods 11, 3297 (2022).

85. Gobbetti, M. et al. Drivers that establish and assembly the lactic acid bacteria biota in cheeses. Trends in Food Science & Technology 78, 244–254 (2018).

86. Bintsis, T. Yeasts in different types of cheese. AIMSMICRO 7, 447–470 (2021).

87. Ceugniez, A., Drider, D., Jacques, P. & Coucheney, F. Yeast diversity in a traditional French cheese “Tomme d’orchies” reveals infrequent and frequent species with associated benefits. Food Microbiology 52, 177–184 (2015).

88. Robertson, P. L. A morphological study of variation in Tyrophagus (Acarina), with particular reference to populations infesting cheese. Bulletin of Entomological Research 52, 501–529 (1961).

89. Aygun, O., Yaman, M. & Durmaz, H. A survey on occurrence of *Tyrophagus putrescentiae* (Acari: Acaridae) in Surk, a traditional Turkish dairy product. Journal of Food Engineering 78, 878–881 (2007).

90. Paillet, T. et al. Virulent phages isolated from a smear-ripened cheese are also detected in reservoirs of the cheese factory. Viruses 14, 1620 (2022).

91. de Melo, A. G., Rousseau, G. M., Tremblay, D. M., Labrie, S. J. & Moineau, S. DNA tandem repeats contribute to the genetic diversity of Brevibacterium aurantiacum phages. Environmental Microbiology 22, 3413–3428 (2020).

92. Kothe, C. I., Bolotin, A., Kraïem, B.-F., Dridi, B. & Renault, P. Unraveling the world of halophilic and halotolerant bacteria in cheese by combining cultural, genomic and metagenomic approaches. International Journal of Food Microbiology 358, 109312 (2021).

93. Juni, E. Psychrobacter. in Bergey’s Manual of Systematics of Archaea and Bacteria 1–10 (John Wiley & Sons, Ltd, 2015). doi:10.1002/9781118960608.gbm01205.

94. Ventosa, A., Haba, R. R., Arahal, D. R. & Sánchez-Porro, C. Halomonas. in Bergey’s Manual of Systematics of Archaea and Bacteria (ed. Oren, A.) 1–111 (Wiley, 2021). doi:10.1002/9781118960608.gbm01190.pub2.

95. Welter, D. K. et al. Free-living, psychrotrophic bacteria of the genus *Psychrobacter* are descendants of pathobionts. mSystems 6, 00258–21 (2021).

96. Saak, C. C. et al. Longitudinal, multi-platform metagenomics yields a high-quality genomic catalog and guides an in vitro model for cheese communities. mSystems 8, e00701–22 (2023).

97. Gilroy, R. et al. Extensive microbial diversity within the chicken gut microbiome revealed by metagenomics and culture. PeerJ 9, e10941 (2021).

98. He, X. et al. Cultivation of a human-associated TM7 phylotype reveals a reduced genome and epibiotic parasitic lifestyle. Proceedings of the National Academy of Sciences 112, 244–249 (2015).

99. Jaffe, A. L., Castelle, C. J. & Banfield, J. F. Habitat transition in the evolution of Bacteria and Archaea. Annual Review of Microbiology 77, 193–212 (2023).

100. VerBerkmoes, N. C., Denef, V. J., Hettich, R. L. & Banfield, J. F. Functional analysis of natural microbial consortia using community proteomics. Nat Rev Microbiol 7, 196–205 (2009).

101. Violle, C. et al. Functional rarity: The ecology of outliers. Trends in Ecology & Evolution 32, 356–367 (2017).

102. de Crécy-lagard, V. et al. A roadmap for the functional annotation of protein families: a community perspective. Database (Oxford) 2022, (2022).

103. Tabuteau, S., Hervé, V., Irlinger, F. & Monnet, C. Metagenomic profiling and genome-centric analysis reveal iron acquisition systems in cheese-associated bacteria and fungi. Environment Microbiology 10.1111/1462-2920.70218 (2026) doi:10.1111/1462-2920.70218.

104. Bonnarme, P., Psoni, L. & Spinnler, H. E. Diversity of L-methionine catabolism pathways in cheese-ripening bacteria. Applied and Environmental Microbiology 66, 5514 (2000).

105. Suganya, K., Govindan, K., Prabha, P. & Murugan, M. An extensive review on L-methioninase and its potential applications. Biocatalysis and Agricultural Biotechnology 12, 104–115 (2017).

106. Abreo, E., Valle, D., González, A. & Altier, N. Control of damping-off in tomato seedlings exerted by *Serratia* spp. strains and identification of inhibitory bacterial volatiles *in vitro*. Systematic and Applied Microbiology 44, 126177 (2021).

107. Inoue, H. et al. Structural analysis of the L-methionine γ-lyase gene from *Pseudomonas putida*. The Journal of Biochemistry 117, 1120–1125 (1995).

108. Alshehri, W. A. Bacterium *Hafnia alvei* secretes L-methioninase enzyme: Optimization of the enzyme secretion conditions. Saudi Journal of Biological Sciences 27, 1222–1227 (2020).

109. Nikulin, A. et al. High-resolution structure of methionine γ-lyase from *Citrobacter freundii*. Acta Crystallographica Section D: Biological Crystallography 64, 211–218 (2008).

110. Malhotra, M., Bello, S. & Gupta, R. S. Phylogenomic and molecular markers based studies on clarifying the evolutionary relationships among *Peptoniphilus* species. Identification of several Genus-Level clades of *Peptoniphilus* species and transfer of some *Peptoniphilus* species to the genus *Aedoeadaptatus*. Systematic and Applied Microbiology 47, 126499 (2024).

111. Suwabe, K., Yoshida, Y., Nagano, K. & Yoshimura, F. Identification of an L-methionine γ-lyase involved in the production of hydrogen sulfide from L-cysteine in *Fusobacterium nucleatum* subsp. *nucleatum* ATCC 25586. Microbiology 157, 2992–3000 (2011).

112. Kothe, C. I. & Renault, P. Metagenomic driven isolation of poorly culturable species in food. Food Microbiology 129, 104722 (2025).

113. Afshari, R., Pillidge, C. J., Dias, D. A., Osborn, A. M. & Gill, H. Cheesomics: the future pathway to understanding cheese flavour and quality. Critical Reviews in Food Science and Nutrition 60, 33–47 (2020).

114. Jin, R. et al. Synthetic microbial communities: Novel strategies to enhance the quality of traditional fermented foods. Comprehensive Reviews in Food Science and Food Safety 23, e13388 (2024).

115. Nikoloudaki, O., Aheto, F., Di Cagno, R. & Gobbetti, M. Synthetic microbial communities: A gateway to understanding resistance, resilience, and functionality in spontaneously fermented food microbiomes. Food Research International 192, 114780 (2024).

116. Jouffret, V. et al. Increasing the power of interpretation for soil metaproteomics data. Microbiome 9, 195 (2021).

