## Supplementary figures and images for "Scratching on French PDO cheese surfaces sheds light on an unexplored microbial genomic and metabolic diversity"

### Suuplementary Figure 1

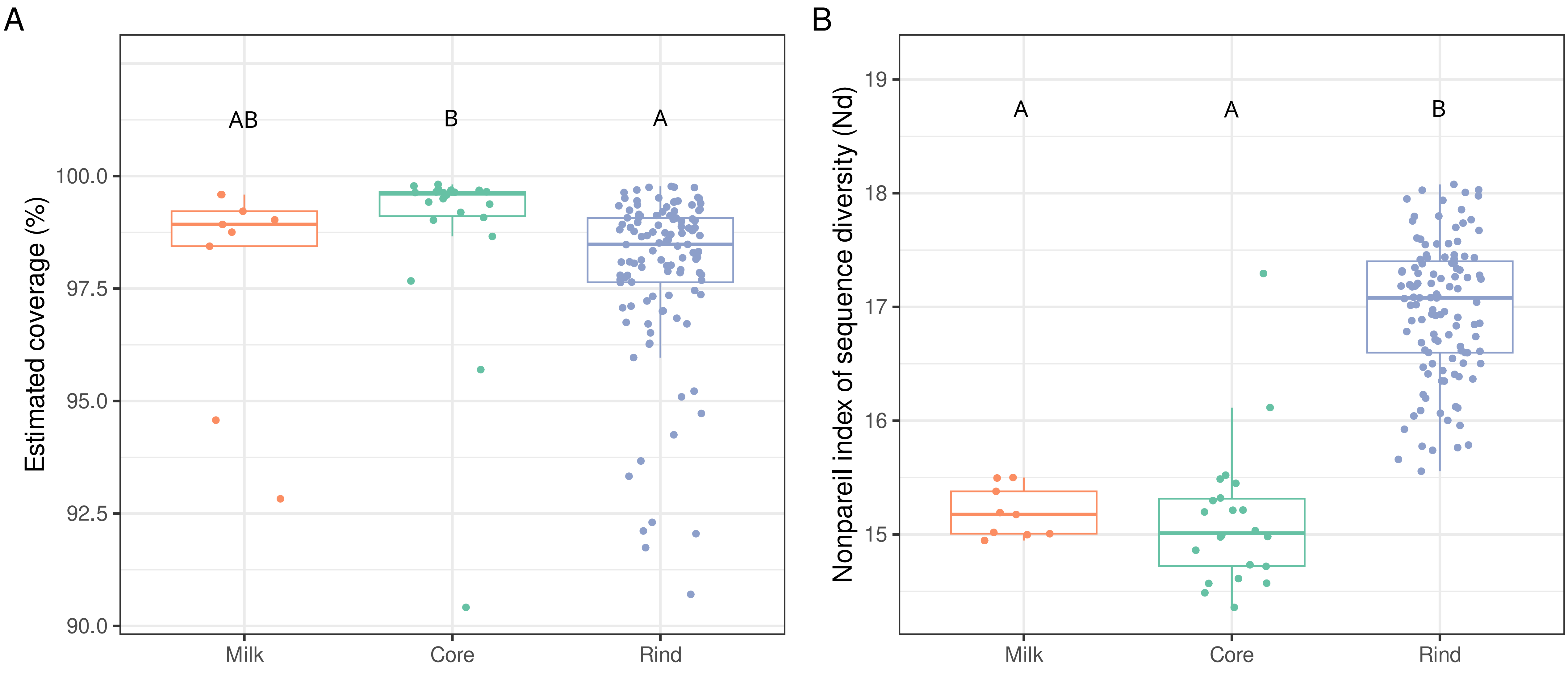

### Suuplementary Figure 2

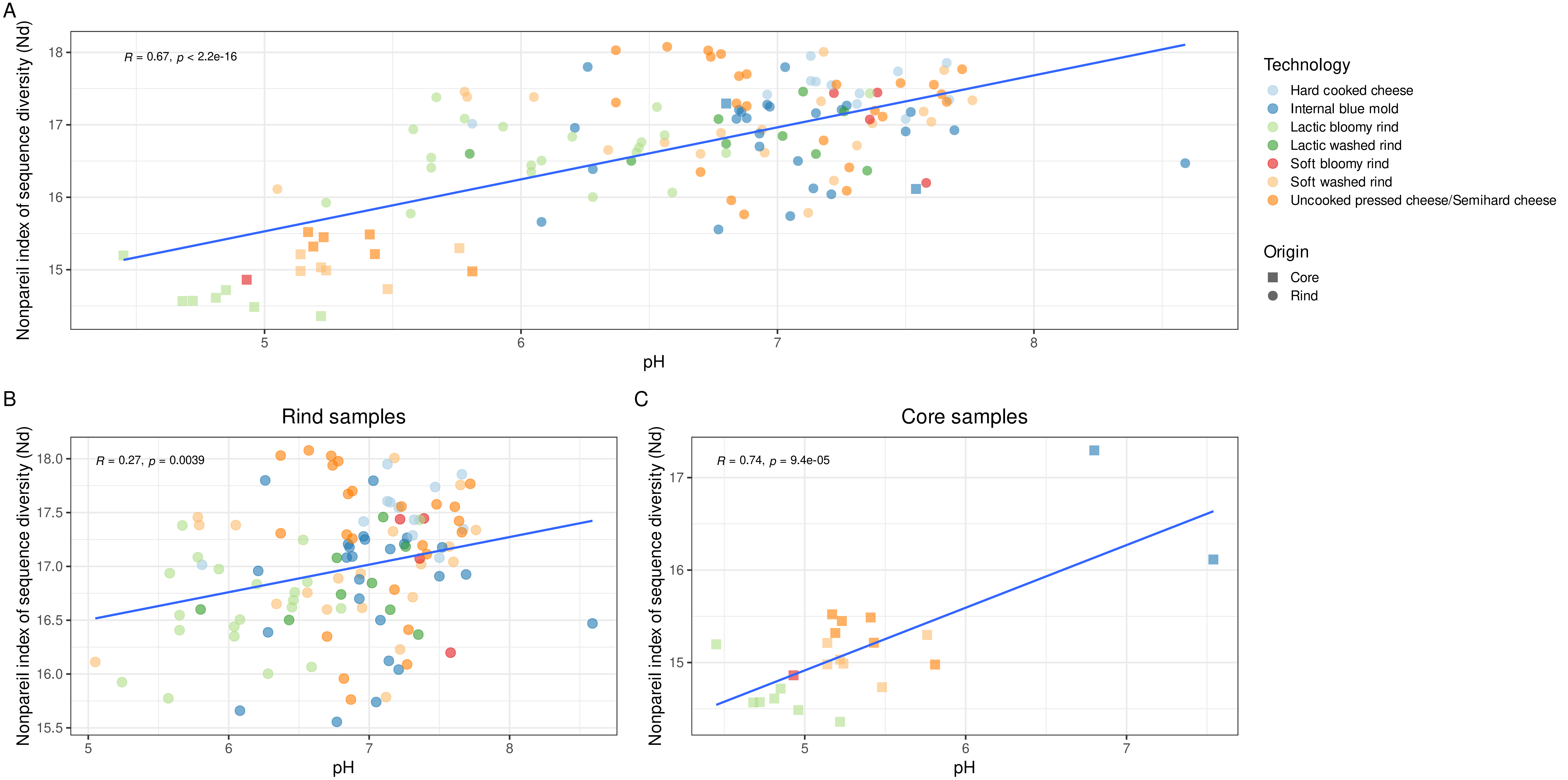

### Suuplementary Figure 3

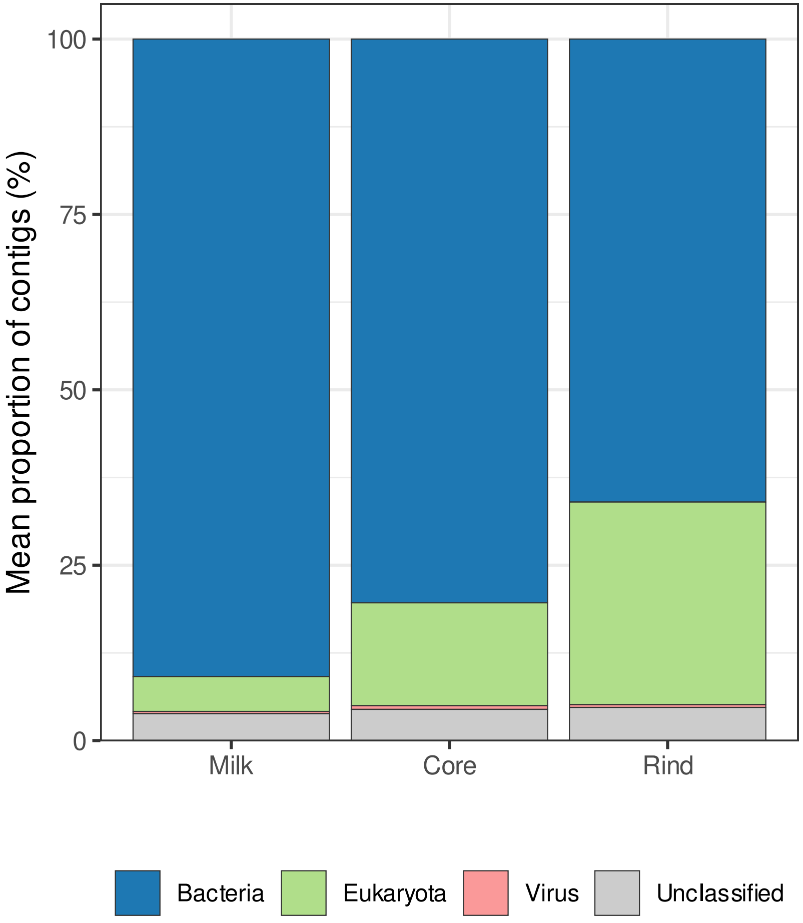

### Suuplementary Figure 4

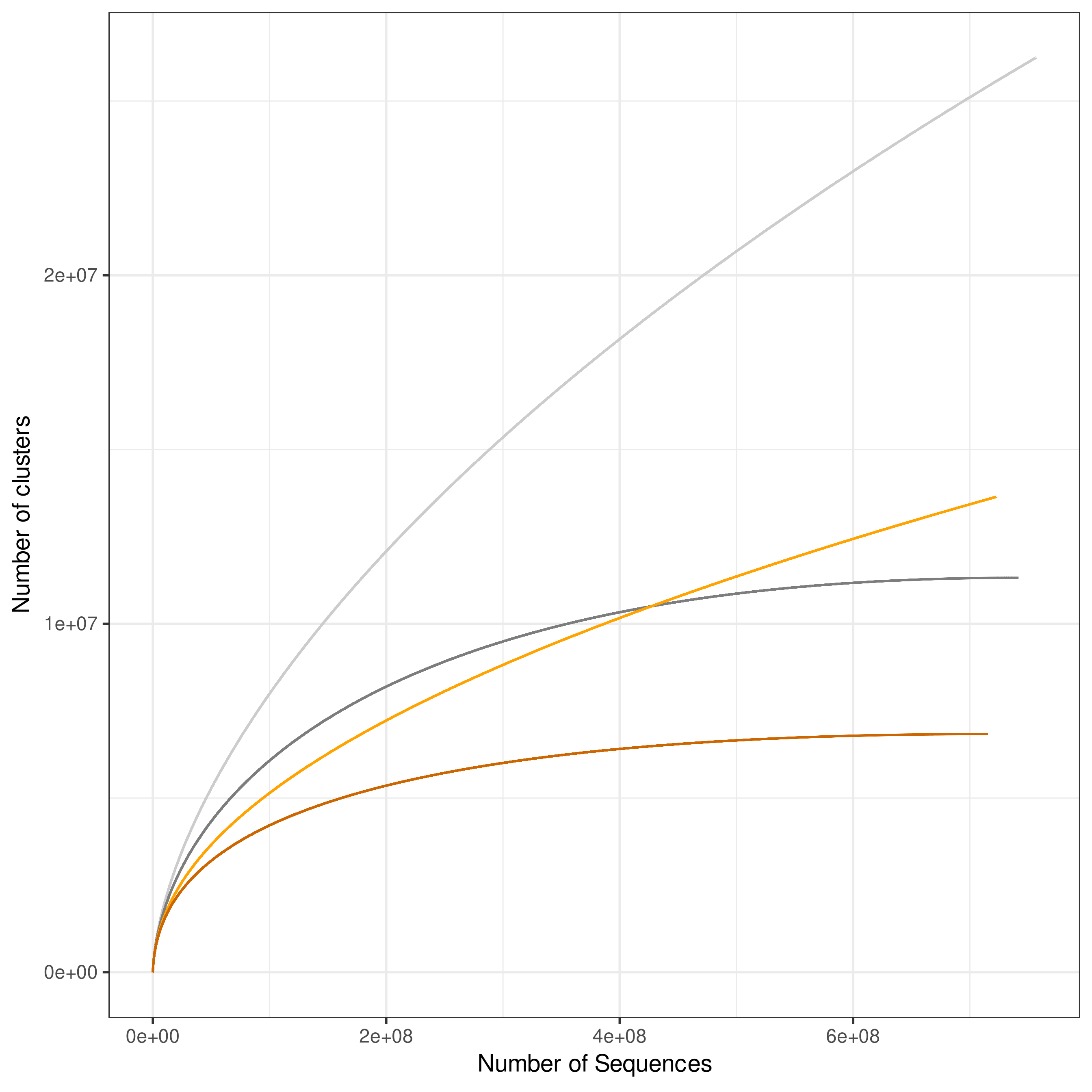

### Suuplementary Figure 5

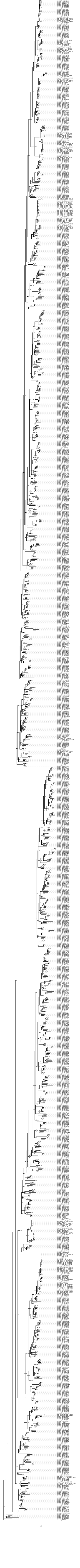
